# The ageing stopping network: Regional and network changes in the IFG, preSMA, and STN across the adult lifespan

**DOI:** 10.1101/2024.08.13.607702

**Authors:** Sarah A. Kemp, Pierre-Louis Bazin, Steven Miletić, Russell J. Boag, Max C. Keuken, Mark R. Hinder, Birte U. Forstmann

## Abstract

Response inhibition, the cancellation of planned movement, is essential for everyday motor control. Extensive fMRI and brain stimulation research provides evidence for the crucial role of a number of cortical and subcortical regions in response inhibition, including the subthalamic nucleus (STN), pre-supplementary motor area (preSMA), and the inferior frontal gyrus (IFG). Current models assume that these regions operate as a network, with action cancellation originating in the cortical areas and then executed rapidly via the subcortex. Response inhibition slows in older age, a change that has been attributed to deterioration or changes in the connectivity and integrity of this network. However, previous research has mainly used whole-brain approaches when investigating changes in structural connectivity across the lifespan, or have used simpler measures to investigate structural ageing. Here, we used high-resolution quantitative and diffusion MRI to extensively examine the anatomical changes that occur in this network across the lifespan. We found substantial changes in iron concentration in these tracts, increases in the apparent diffusion coefficient, and some evidence for demyelination. Conversely, we found very little evidence for age-related anatomical changes in the regions themselves. We propose that some of the functional changes observed in these regions in older adult populations (e.g., increased BOLD recruitment) are a reflection of alterations to the connectivity between the regions, rather than localised regional change.

## 1 Introduction

Response inhibition is an essential part of everyday motor control, and is crucial for successfully navigating dynamic and complex environments. An important part of response inhibition is action cancellation, the sudden termination of planned or already-initiated movement (Sebastian, Pohl, et al., 2013; Wessel & Aron, 2017). Action cancellation is impeded in older age, with older adults taking longer to cancel their movements (Rey-Mermet and Gade, 2018; Smittenaar et al., 2013; Tsvetanov et al., 2018). This change has been attributed both to neural breakdown in the regions responsible for action cancellation and changes in response inhibition strategies in older adults (Bissett & Logan, 2011; Bloemendaal et al., 2016; van de Laar et al., 2011). The specific neural breakdown that occurs in these cortical and subcortical regions is incompletely understood. Here, we combine multiple imaging approaches to extensively examine the anatomical changes that occur in this network across the adult lifespan.

The neural correlates of response inhibition and action cancellation are highly complex, and can vary depending on the specificities of the task (Isherwood, Bazin, et al., 2023; Puri et al., 2018; Sebastian, Pohl, et al., 2013). However, a large body of brain stimulation and imaging research has consistently identified the subthalamic nucleus (STN), inferior frontal gyrus (IFG), and pre-supplementary motor area (preSMA) as being intrinsically involved in action cancellation (see e.g., Aron et al., 2007; Isherwood, Kemp, et al., 2023; Isherwood et al., 2021; Lee et al., 2016; Lofredi et al., 2021; Xu et al., 2016; Zandbelt et al., 2013). Current models of response inhibition propose that these regions form a complex ’stopping network’, and regulate the initiation and cancellation of actions through three cortico-basal ganglia pathways (the direct, indirect, and hyperdirect pathways), which send commands to the effector muscles via the STN and other parts of basal ganglia (Graybiel, 2000; Rocha et al., 2023; Schroll & Hamker, 2013). This framework was initially developed from research involving rodents and non-human primates (see e.g., Coudé et al., 2018; Eagle et al., 2008; Nambu et al., 2002; Nougaret et al., 2013; S. Schmidt et al., 2009), but the anatomical plausibility of the hyperdirect pathway in humans has recently been established (Chen et al., 2020). Action cancellation itself was originally thought to be implemented via the hyperdirect pathway, originating in the IFG (Aron & Poldrack, 2006; Aron et al., 2004, 2014; Bingham et al., 2023; Cavanagh et al., 2014), while two-stage models of action cancellation hypothesise that both the indirect and hyperdirect pathways are critically involved in action cancellation, originating in humans in the preSMA and IFG, respectively (Diesburg & Wessel, 2021; Frank, 2006; R. Schmidt & Berke, 2017). As well as their individual contributions to action cancellation, the *connectivity* of these three regions has been shown to be an important predictor of stopping performance and other measures of response inhibition (Boen et al., 2022; Forstmann et al., 2012; King et al., 2012; Rae et al., 2015; Singh et al., 2021). This highlights the importance of considering how these regions cooperate and interact as a network in order to effectively coordinate movement, as well as considering their individual roles.

Action cancellation is frequently investigated using the stop-signal task (SST), where participants have to make a default "go" response in every trial, occasionally needing to cancel or inhibit this response after it has been initiated. Performance in trials where the response is cancelled is normally quantified by the stop-signal reaction time (SSRT), which estimates the speed of the latent stop process (Verbruggen et al., 2019). Older adults generally exhibit slower response times in SSTs and other response inhibition tasks (see e.g., Healey et al., 2024; Kang et al., 2022; Nikitenko et al., 2020; Rey-Mermet and Gade, 2018), but also show notably longer SSRTs compared to younger adults (Hsieh & Lin, 2017; van de Laar et al., 2011; Williams et al., 1999). Importantly, this increase in SSRT cannot solely be attributed to the aforementioned general slowing of response times (Bedard et al., 2002; Hu et al., 2018), indicating there is neural degradation unique to the stopping process in older age.

Alongside these behavioural changes, older adults also show changed neural recruitment patterns in response inhibition tasks. During SSTs, older adults tend to show less activation in the traditional ’stopping regions’ (STN, preSMA, IFG), but increased activity in a wide range of additional regions (Hu et al., 2019; Hu et al., 2018; Kang et al., 2022; Kleerekooper et al., 2016; Sebastian, Baldermann, et al., 2013). These broader recruitment patterns have also been observed in other tasks (see e.g., Kennedy et al., 2015; Zhu et al., 2010). Theoretical frameworks such as the dedifferentiation hypothesis, Scaffolding Theory of Ageing and Cognition (STAC), and Compensation-Related Utilisation of Neural Circuits Hypothesis (CRUNCH) postulate that these broader recruitment patterns serve as a compensatory mechanism for localised deterioration in the regions that were previously more specialised in performing these tasks (Geerligs et al., 2015; Kang et al., 2022; Reuter-Lorenz & Park, 2014). This more widespread recruitment serves to maintain behavioural performance, but as task demand increases or when there is further structural degradation, this compensation is less effective and behavioural performance begins to decline. Age-related structural decline in the brain has indeed been observed, but patterns of decline vary from region to region and between different networks (Geerligs et al., 2015; Zimmermann et al., 2016). Changes in specific networks and regions need therefore to be individually considered and cannot be described in a generalised manner.

Age-related connectivity changes have previously been investigated using diffusion-weighted imaging (DWI), a structural application of magnetic resonance imaging (MRI) which quantifies the movement of water molecules in the brain. This movement is anatomically constrained by brain structures such as white matter and fibre bundles. Determining the magnitude and direction of water diffusion thus enables estimation of the underlying anatomy (Grier, 2020; O’Donnell & Westin, 2011). There are a number of common diffusion metrics, but two widely-used ones are the apparent diffusion coefficient (ADC) and fractional anisotropy (FA) (Beck et al., 2021; Lazari & Lipp, 2021; Porcu et al., 2021; Sullivan & Pfefferbaum, 2006). The ADC relates to the net diffusion in a voxel, and will increase as water movement becomes less anatomically constrained (e.g., with breakdown of brain structure). FA is a ratio measure that quantifies the extent to which the water movement is anisotropic (non-uniform). Higher values of FA (closer to 1) indicate that the movement is more constrained. Both measures are generally thought to reflect white matter integrity and be influenced by biophysical measures (Grier, 2020; O’Donnell & Westin, 2011). These measures have frequently been used to investigate neuroanatomical changes across the adult lifespan (see e.g., Beck et al., 2021; Henriques et al., 2023; Lebel et al., 2012). The changes that occur in these metrics in older age vary from region to region, but generally, FA will increase and ADC will decrease

Connectivity between the canonical stopping regions (STN, IFG, preSMA), indexed via various diffusion metrics, has been associated with response inhibition performance and SSRT in children, adolescents, and young adults (Aron, 2007; Boen et al., 2022; King et al., 2012; Madsen et al., 2020; F. Zhang & Iwaki, 2020). While there has been less investigation of older adults, broader changes in diffusion indices (i.e., not specific to the stopping network) have been linked to response inhibition performance (Yang et al., 2019) and connectivity between the preSMA and STN has been found to be predictive of SSRT (Coxon et al., 2012). Aside from these few examples, investigations regarding the anatomical changes that occur in this network across the adult lifespan have been limited, particularly in healthy populations.

Further, although the aforementioned diffusion metrics are often assumed to directly index myelin integrity, they are likely influenced by a range of microstructural properties in older adults, including axonal loss, fibre dispersion, and variability in fibre and neurite orientation (Beck et al., 2021; Grussu et al., 2017; Lazari & Lipp, 2021), and are thus relatively unspecific in terms of the anatomical changes they capture. Assessments of neuroanatomical ageing can therefore be complemented by other imaging techniques. Quantitative MRI (qMRI) can be used to estimate histological measures of brain tissue, such as iron and myelin, *in vivo* (Keuken et al., 2017; Madden & Merenstein, 2023; Miletić et al., 2022; Weiskopf et al., 2015). As well as being critical for normal brain function, these biophysical properties are highly relevant in investigations of the ageing brain. Demyelination is a hallmark of ageing, and iron is increasingly recognised as an important factor in both diseased and healthy ageing contexts (Buyanova & Arsalidou, 2021). Importantly, age-related changes in iron and myelin concentrations tend to be region-specific, with certain areas exhibiting high levels of iron accumulation and others showing minimal changes (Daugherty & Raz, 2015; Hagemeier et al., 2012; Miletić et al., 2022). Taken together, assessment and interpretation of neuroanatomical changes across the lifespan needs to be quite region-specific, i.e., generalised hypotheses may not apply to individual networks or regions (Geerligs et al., 2015). Utilisation of multimodal imaging approaches aids to build a richer picture of anatomical change, enhancing understanding of the intricate and dynamic changes that occur across the adult lifespan (Tardif et al., 2016).

Here, we combined high-resolution DWI and ultra-high field qMRI (acquired at 3 and 7 Tesla, respectively) to examine the physiology of the stopping network across the lifespan in an adult sample. We quantified age-related changes in these measures, providing a comprehensive assessment of the neuroanatomical changes that may precipitate the behavioural changes in action cancellation observed in older populations. Additionally, given that diffusion measures are so frequently assumed to index myelination and the limited availability of DWI and qMRI datasets in the same sample, we explored the relationship between the diffusion measures and age, iron, and myelin to guide the interpretation of DWI metrics in future studies.

## 2 Methods

### 2.1 Participants

We used the Amsterdam ultra-high field adult lifespan database (AHEAD; Alkemade et al., 2020), which comprises multimodal MRI data from 105 healthy adults. We used the subset of these (*N* = 49, 27 females) for whom diffusion-weighted imaging (DWI) data are also available (Keuken et al., 2022). Participants were scanned on two different days, some weeks apart. The quantitative images were acquired in the first session and the diffusion images were acquired in the second. The overarching procedure is depicted in Figure 1. Participants were aged between 18-90 years old. Exclusion criteria were any factors that may interfere with MRI scanning, e.g., MRI incompatibility, pregnancy, or self-reported claustrophobia. All procedures were in accordance with the Code of Ethics of the World Medical Association and approved by the Institutional Review Board at the University of Amsterdam (ERB number 2016-DP-6897). Informed consent was obtained from participants including permission for the future release of de-identified data.

**Figure 1:**
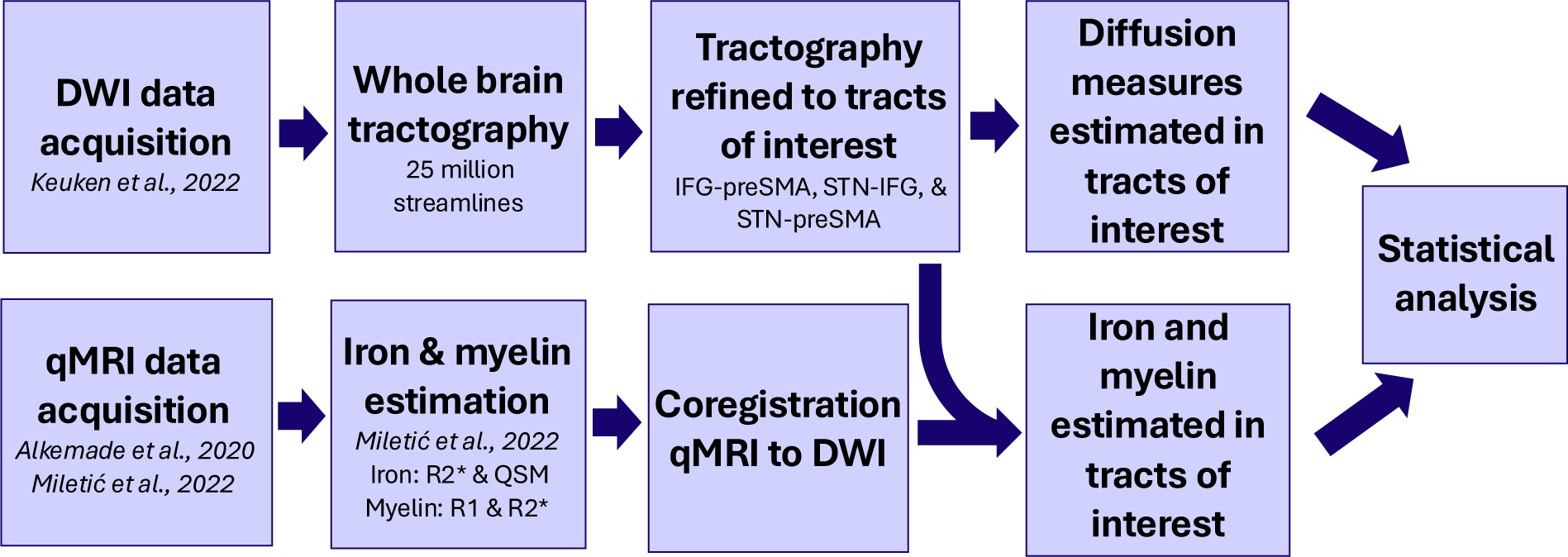
Image acquisition and processing steps. Participants were scanned on two different days, with the qMRI images acquired in the first session and the DWI images in the second. All processing was initially conducted in individual space, and then ultimately co-registered to MNI space.

### 2.2 Diffusion-weighted imaging (DWI)

#### 2.2.1 Data acquisition

The diffusion images were acquired at the Spinoza Centre for Neuroimaging in Amsterdam, the Netherlands with a Philips 3 T Ingenia CX MRI scanner using a 32-channel receiver head coil. We used an MS-ME (multi slice spin-echo) single shot sequence with 1.28 mm isotropic resolution. Diffusion weighting was isotopically distributed along 48, 56, or 64 directions with b-values of 700 s/mm^2^, 1000 s/mm^2^, and 1600 s/mm^2^, respectively. Due to memory limitations, the 64-direction scan was split into two 32-direction scans. Twelve b0 volumes were acquired within the MS-ME sequences and an additional eight were acquired at the end with an inverted fat shift direction. Scan parameters: TE = 78 ms, FA = 90°, TR = 8100 ms, acceleration factor SENSE_AP_ = 2, FOV = 205 × 205, slice gap = 0 mm, TA_b0_ = 00:49 min, TA_b700_ = 13:48 min, TA_b1000_ = 17:19 min, TA_b1600_ = 9:44 min for each half, i.e., 19:28 min in total. Total acquisition time ≈ 100 minutes.

#### 2.2.2 Preprocessing

The preprocessing pipeline for this diffusion dataset has been previously reported (Keuken et al., 2022). In brief, all data were denoised using the PCA-based function *dwidenoise* in MRtrix V3.0.2 (Tournier et al., 2019). Gibbs artefacts were removed using the *mrdegibbs* function in MRtrix, and susceptibility-induced off-resonance field distortions were corrected using the different phase-encoding blips of the b0 images in *topup*, implemented with FSL V5.0.11 (S. M. Smith et al., 2004). The output from *topup* was used to apply eddy current correction using *eddyOpenMP* in FSL. Finally, an N4 bias correction was applied using *dwibiascorrect* and a brain mask estimated using *dwi2mask*, both with MRtrix.

#### 2.2.3 Tractography

Connectivity between the three ROIs (IFG-preSMA, IFG-STN, and STN-preSMA in each hemisphere) was estimated from whole-brain tractography, implemented using MRtrix (Tournier et al., 2019). This process was performed in individual space for each participant, and the final six tracts were transformed into MNI space.

From the preprocessed diffusion images (see 2.2.2), fibre orientation distributions (FODs) were estimated using *dwi2fod* (Tournier et al., 2004). We used the *msmt_csd* algorithm, which is designed for data with multiple b-values (Jeurissen et al., 2014). We then performed wholebrain tractography based on the FODs using *tckgen* with 25 million streamlines. Here we used the iFOD2 (Second-Order Integration Over FODs) algorithm, a probabilistic algorithm that is more optimised for regions of high curvature (Tournier et al., 2010). Note that we adjusted the FOD amplitude cutoff point for track termination from 0.1 to 0.05; we found the original cutoff value to be too conservative for some of the older adults in the dataset, drastically limiting the number of streamlines identified for those participants in regions with white matter inflammation or lesions.

We then refined the whole-brain tractograms using *tcksift2* (R. E. Smith et al., 2015) and isolated the streamlines that were connected to any of the ROIs using *tck2connectome* (Hagmann et al., 2008). The STN was defined based on the Multi-Contrast Anatomical Subcortical Structures Parcellation (MASSP) automated algorithm (Bazin et al., 2020). The IFG and preSMA were defined based on masks from Neubert and colleagues (IFG: Neubert et al., 2014; preSMA: Neubert et al., 2015). Subsampled tractograms from two participants are shown in Figure 2.

**Figure 2:**
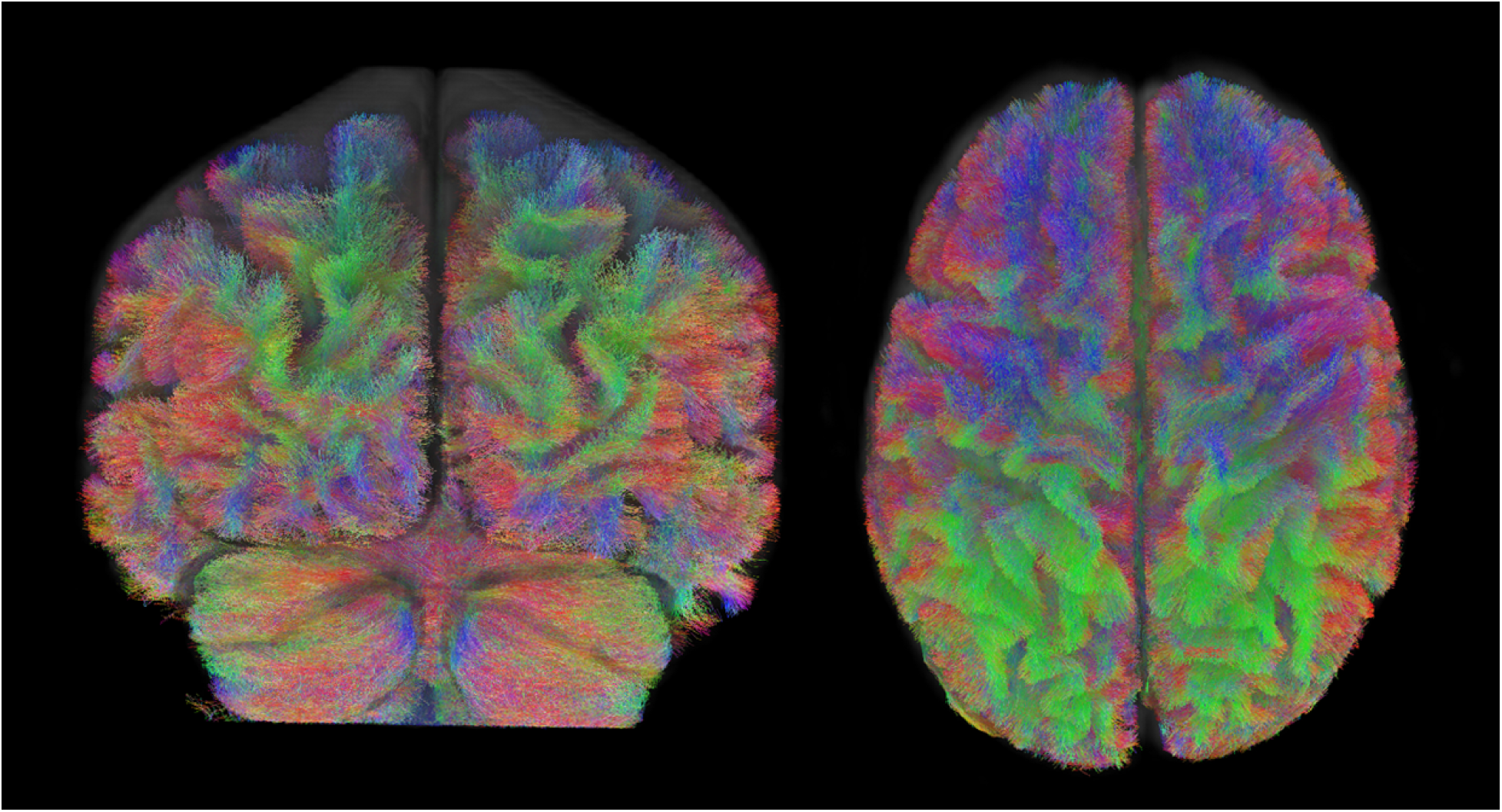
Whole-brain tractograms from two random but representative participants, shown in the coronal (left) and axial (right) planes. Note that these images show subsampled tractograms, with 200 000 streamlines. The tractography used for analysis had 25 million streamlines, estimated using fibre orientation distributions.

Once we had the refined tractogram from *tck2connectome*, we used *connectome2tck* to identify the streamlines between each pair of the three ROIs, i.e., the streamlines between IFG-preSMA, STN-IFG, and STN-preSMA in each hemisphere. An example of three of the six tracts for one participant can be found in Figure 3.

**Figure 3:**
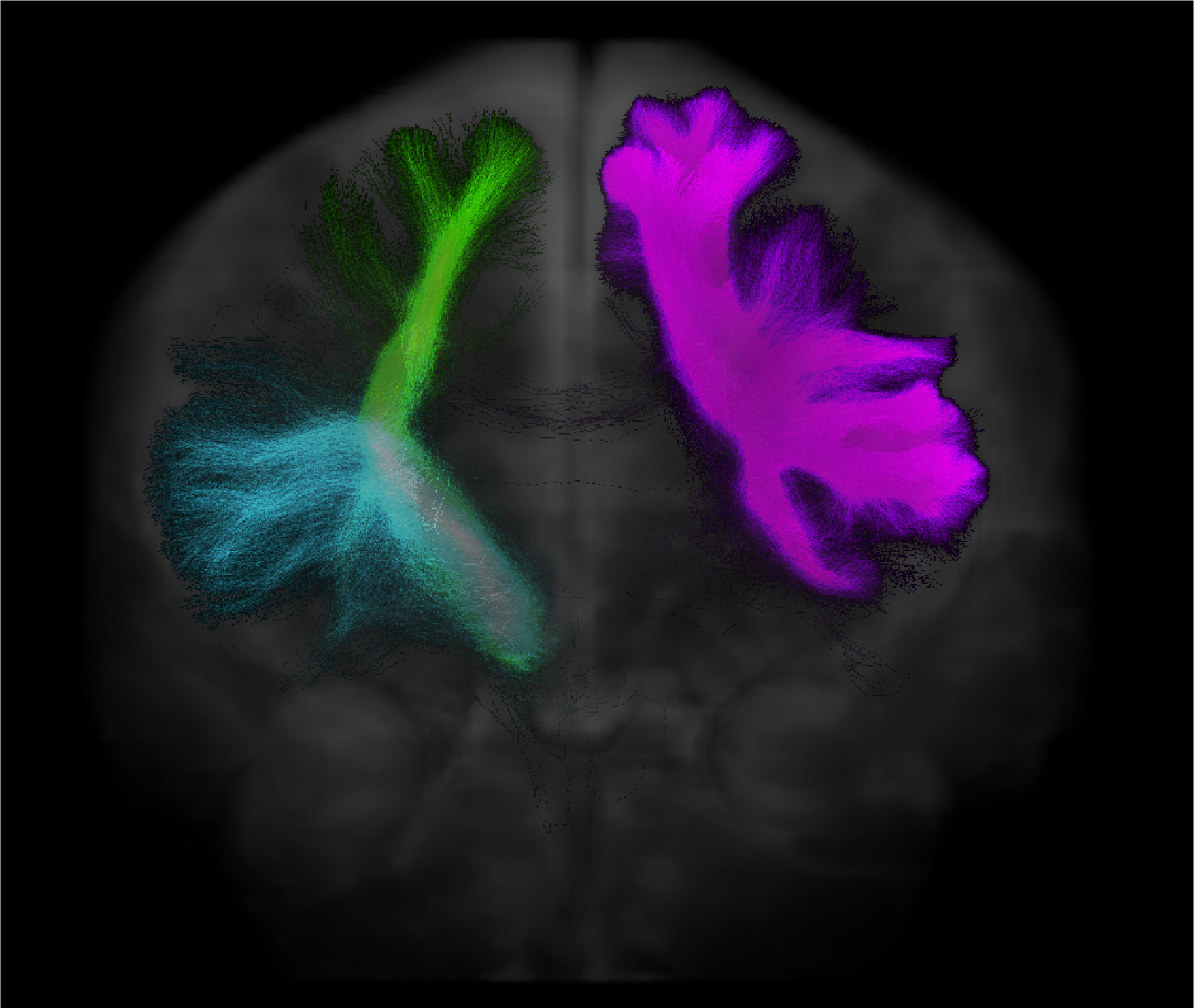
Three of the six tracts of interest from one participant. The tracts of interest were the STN-IFG, STN-preSMA, and IFG-preSMA in each hemisphere, but for ease of visualisation, each tract is only shown in one hemisphere here. The depicted tracts are right STN-IFG (blue), right STN-preSMA (green), and left IFG-preSMA (pink), imposed over the mean b0 image.

#### 2.2.4 Estimation of generalised FA and ADC

Rather than the conventional estimation methods for FA, using the diffusion tensor model, we estimated generalised FA (GFA) as per Tuch (2004):

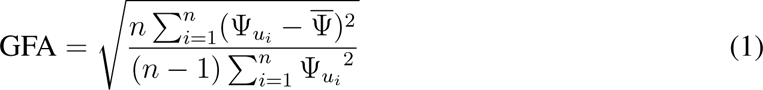

This approach makes use of the white matter FODs (generated in 2.2.3), with Ψ given as the FOD coefficients. Usage of the FODs in this manner retains multidirectional anisotropy information, making this estimation method more useful in places of high curvature, or with a particular abundance of crossing fibres (Tan et al., 2015; Tuch, 2004). It should be interpreted in the same manner as standard FA (i.e., it quantifies the extent to which water movement is anisotropic in that voxel).

ADC was calculated for each of the three b-values, and these values were averaged to get a voxelwise, whole-brain estimation of ADC per participant (Moreau et al., 2018):

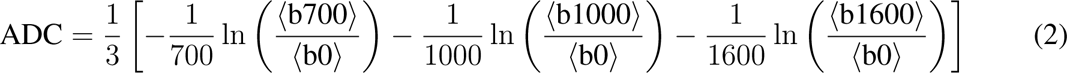

### 2.3 Quantitative MRI (QMRI)

#### 2.3.1 Data acquisition

The quantitative images were acquired at the Spinoza Centre for Neuroimaging with a Philips 7 T Achieva MRI scanner with a 32-channel, phased-array coil. T_1_, T_2_*, and QSM contrasts were obtained using an MP2RAGEME (multi-echo magnetisation-prepared rapid gradient echo) sequence (Caan et al., 2019) with 0.7 mm isotropic resolution. MP2RAGEME is an extension of the MP2RAGE sequence (Marques et al., 2010), and acquires two rapid gradient echo images (GRE1 and GRE2) in the sagittal plane after a 180-degree inversion pulse and excitation pulses with inversion times of TI_1_ = 670 ms and TI_2_ = 3675.4 ms. A multi-echo readout was added to the second inversion at four echo times (TE_1_ = 3 ms, TE_2,1−4_ = 3, 11.5, 19, 28.5 ms). Other scan parameters: FA_GRE1,GRE2_ = (4°, 4°), TR_GRE1_ = 6.2 ms, TR_GRE2_ = 31 ms, TR_MPRAGEME_ = 6778 ms, acceleration factor SENSE_PA_ = 2, FOV = 205 × 205 × 164 mm, acquisition matrix = 292 × 290, bandwidth = 404.9 MHz, reconstructed voxel size = 0.67 × 0.67 × 0.7 mm, TFE (turbo factor) = 150 resulting in 176 shots. Total acquisition time = 19.53 min.

#### 2.3.2 Iron and myelin approximation

The procedure for estimating iron and myelin from this sample has been reported in full elsewhere (Miletić et al., 2022). In brief, preprocessing and the reconstruction of quantitative values was implemented in Python using Nighres (Huntenburg et al., 2018). Before reconstruction, the raw images were skull-stripped and denoised using LCPCA (Bazin et al., 2019). Voxelwise values of iron and myelin were predicted for each participant based on R1, R2*, and quantitative susceptibility mapping (QSM) metrics as follows:

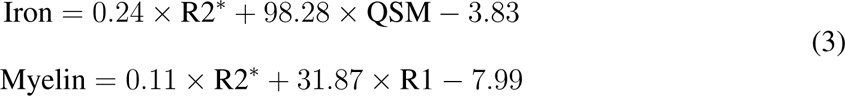

The models for iron and myelin approximation were each selected from a series of eight linear models, fitted with all possible combinations of R1, R2*, and QSM using ordinary least squares (OLS). These estimations yielded whole-brain maps with iron and myelin estimates per voxel for each participant. From here, we applied the masks for each of the ROIs and for the tracts of interest generated in 2.2.3, resulting in estimations of iron and myelin concentration in those regions and tracts for each participant. The overarching procedure is depicted in Figure 1.

### 2.4 Data analysis

We investigated age-related regional and network changes in the stopping network across four measures: iron, myelin, GFA, and ADC. We first assessed age-related alterations in iron and myelin in the IFG, preSMA, and STN (regional changes), and then in iron, myelin, GFA, and ADC in the IFG-preSMA, STN-IFG, and STN-preSMA (network changes).

In each instance, we fit a series of polynomial regression models with participant age as a linear, quadratic, and/or cubic term. All models were fit with OLS, implemented using the Python library *statsmodels* (Seabold & Perktold, 2010). Given the varying number of parameters in our candidate models, we compared the relative model fits using Bayesian information criterion (BIC; Schwarz, 1978). In each case, the model with the lowest BIC was taken as the winning model. For each winning model, we then identified influential datapoints using Cook’s distance, with a threshold cutoff of 4/*n* (≈ 0.08; Rawlings et al., 1998). Individual datapoints that exceeded this threshold were removed. We then refit the data to each of the candidate models and re-selected the winning model, again using BIC for model comparison.

Finally, we further quantified the relationship between age and the diffusion measures using a mediation analysis (again using *statsmodels*; Imai et al., 2010; Seabold and Perktold, 2010), with iron and myelin given as possible mediators.

## 3 Results

### 3.1 Averaging between hemispheres and outlier analysis

Student *t*-tests were used to assess if there were any significant differences between hemispheres for any of the outcome variables for any region or tract. All *t*-tests were non-significant (all *p > .*400). We therefore collapsed the data across hemispheres to reduce the number of fitted models and increase statistical power. The results of all *t*-tests can be found in the Supplementary Materials.

Outliers were identified using Cook’s distance. After identifying each winning model, we removed influential datapoints using a threshold cutoff of 4/*n* (Rawlings et al., 1998) and refitted all candidate models on the data, excluding these datapoints, and re-selecting the winning model. This removed an average of 2.67 datapoints per model (5.44% of total datapoints). Exclusions for each model can be found in the analysis code (https://osf.io/hnz79/).

### 3.2 Regional changes (IFG, preSMA, STN)

We fit a series of polynomial regressions to assess age-related changes in each of the individual regions of interest (IFG, preSMA, STN) for iron and myelin, with participant age as a linear, quadratic, and/or cubic term, using BIC for model comparison. The six model candidates can be found in the Supplementary Materials. Model comparison results for all candidate models (including BIC values) can be found at https://osf.io/hnz79/.

Parameterised winning models with corresponding *R*^2^ are shown in Table 1. Winning models showed linear age-related changes in myelin for all three ROIs. Age-related changes in iron were found to be linear for the IFG and preSMA, and quadratic for the STN.

**Table 1:** Age-related changes in iron and myelin in the IFG, preSMA, and STN.

| Region | Measure | Winning model | $R^2$ |
| --- | --- | --- | --- |
| IFG | Iron | $\gamma = 3.858 + 0.015x$ | 0.155 |
| | Myelin | $\gamma = 7.549 - 0.009x$ | 0.038 |
| preSMA | Iron | $\gamma = 4.510 - 0.010x$ | 0.071 |
| | Myelin | $\gamma = 8.046 + 0.031x$ | 0.059 |
| STN | Iron | $\gamma = 6.732 + 0.371x - 0.004x^2$ | 0.080 |
| | Myelin | $\gamma = 15.353 - 0.011x$ | 0.025 |

Parameterised winning models showing age-related changes in iron and myelin in the regions of interest. $\gamma$ denotes the outcome variable (iron or myelin); $x$ denotes age. Winning models were selected based on BIC.

### 3.3 Network changes (IFG-preSMA, STN-IFG, STN-preSMA)

#### 3.3.1 Age-related changes in iron, myelin, GFA, and ADC

We first assessed age-related changes in the stopping network for the biophysical (iron and myelin) and diffusion measures (GFA and ADC). As in 3.2, we fit a series of polynomial regression models with participant age as a linear, quadratic, and/or cubic term, using BIC for model comparison. See Supplementary Materials for all candidate models. Model comparison results for all candidate models (including BIC values) can be found at https://osf.io/hnz79/. The parameterised winning models are described below and reported in Table 2 with their corresponding *R*^2^.

**Table 2:**
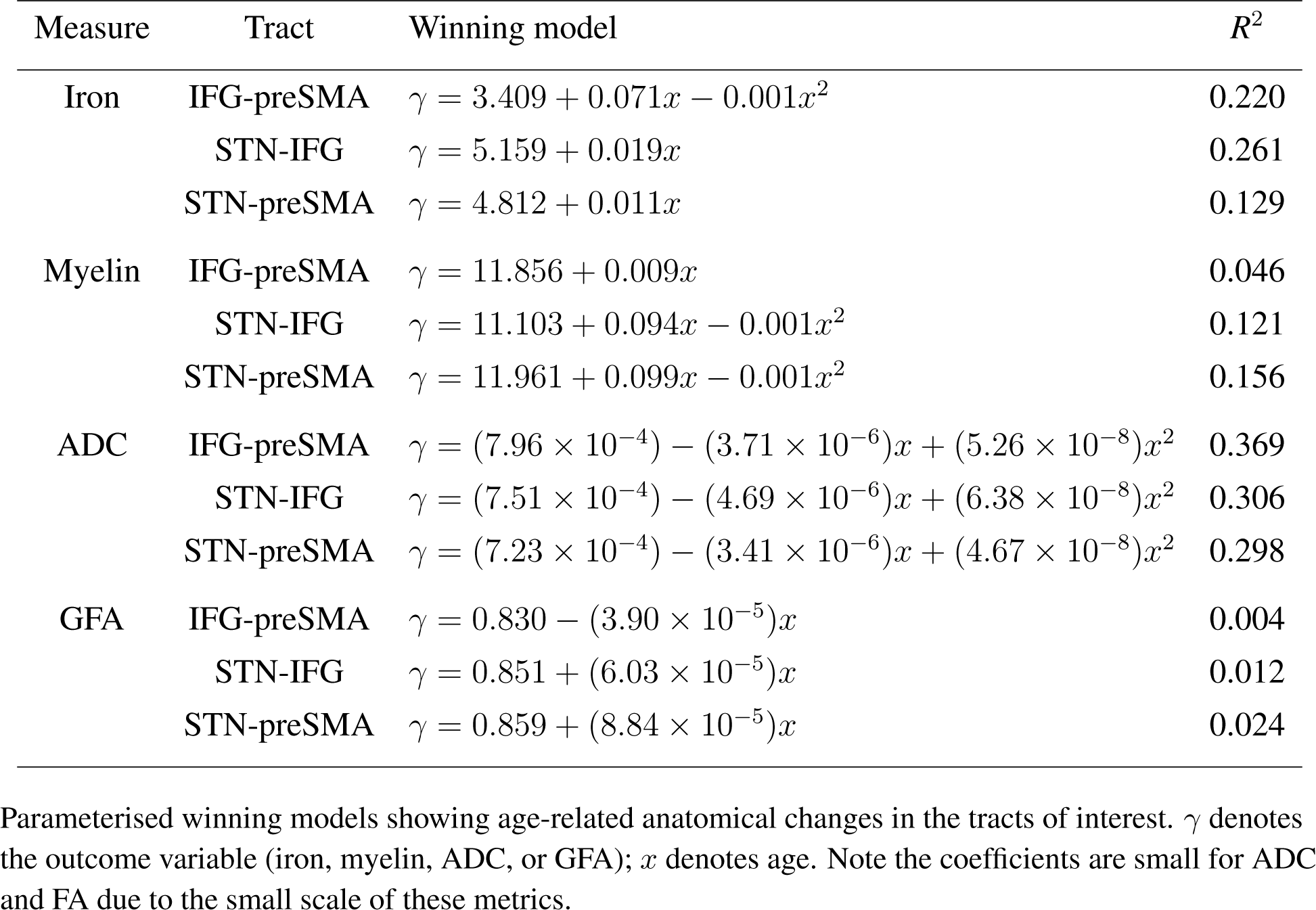
Age-related changes in the stopping network.

| Measure | Tract | Winning model | $R^2$ |
| --- | --- | --- | --- |
| Iron | IFG-preSMA | $\gamma = 3.409 + 0.071x - 0.001x^2$ | 0.220 |
| | STN-IFG | $\gamma = 5.159 + 0.019x$ | 0.261 |
| | STN-preSMA | $\gamma = 4.812 + 0.011x$ | 0.129 |
| Myelin | IFG-preSMA | $\gamma = 11.856 + 0.009x$ | 0.046 |
| | STN-IFG | $\gamma = 11.103 + 0.094x - 0.001x^2$ | 0.121 |
| | STN-preSMA | $\gamma = 11.961 + 0.099x - 0.001x^2$ | 0.156 |
| ADC | IFG-preSMA | $\gamma = (7.96 \times 10^{-4}) - (3.71 \times 10^{-6})x + (5.26 \times 10^{-8})x^2$ | 0.369 |
| | STN-IFG | $\gamma = (7.51 \times 10^{-4}) - (4.69 \times 10^{-6})x + (6.38 \times 10^{-8})x^2$ | 0.306 |
| | STN-preSMA | $\gamma = (7.23 \times 10^{-4}) - (3.41 \times 10^{-6})x + (4.67 \times 10^{-8})x^2$ | 0.298 |
| GFA | IFG-preSMA | $\gamma = 0.830 - (3.90 \times 10^{-5})x$ | 0.004 |
| | STN-IFG | $\gamma = 0.851 + (6.03 \times 10^{-5})x$ | 0.012 |
| | STN-preSMA | $\gamma = 0.859 + (8.84 \times 10^{-5})x$ | 0.024 |

Parameterised winning models showing age-related anatomical changes in the tracts of interest. $\gamma$ denotes the outcome variable (iron, myelin, ADC, or GFA); $x$ denotes age. Note the coefficients are small for ADC and FA due to the small scale of these metrics.

### Iron and myelin

The winning models showed linear age-related increases in iron in the STN-IFG and STN-preSMA tracts, and quadratic changes in the IFG-preSMA tract. The winning models showed quadratic changes in myelin in the STN-IFG and STN-preSMA tracts, and linear changes in the IFG-preSMA tract.

#### ADC and GFA

The winnings models identified quadratic changes in ADC for all three tracts of interest and linear changes for GFA. The coefficients for these models are particularly small due to their scale; normal white matter ADC ranges from 0.60 − 1.05 × 10^−3^mm^2^/s (Sener, 2001), while GFA is a ratio, i.e., values are between 0 and 1 (Figley et al., 2022). Note also that *R*^2^ < 0.02 for all three winning models for GFA, and should be interpreted with caution.

Figure 4 shows the winning models for iron, myelin, ADC, and GFA.

**Figure 4:**
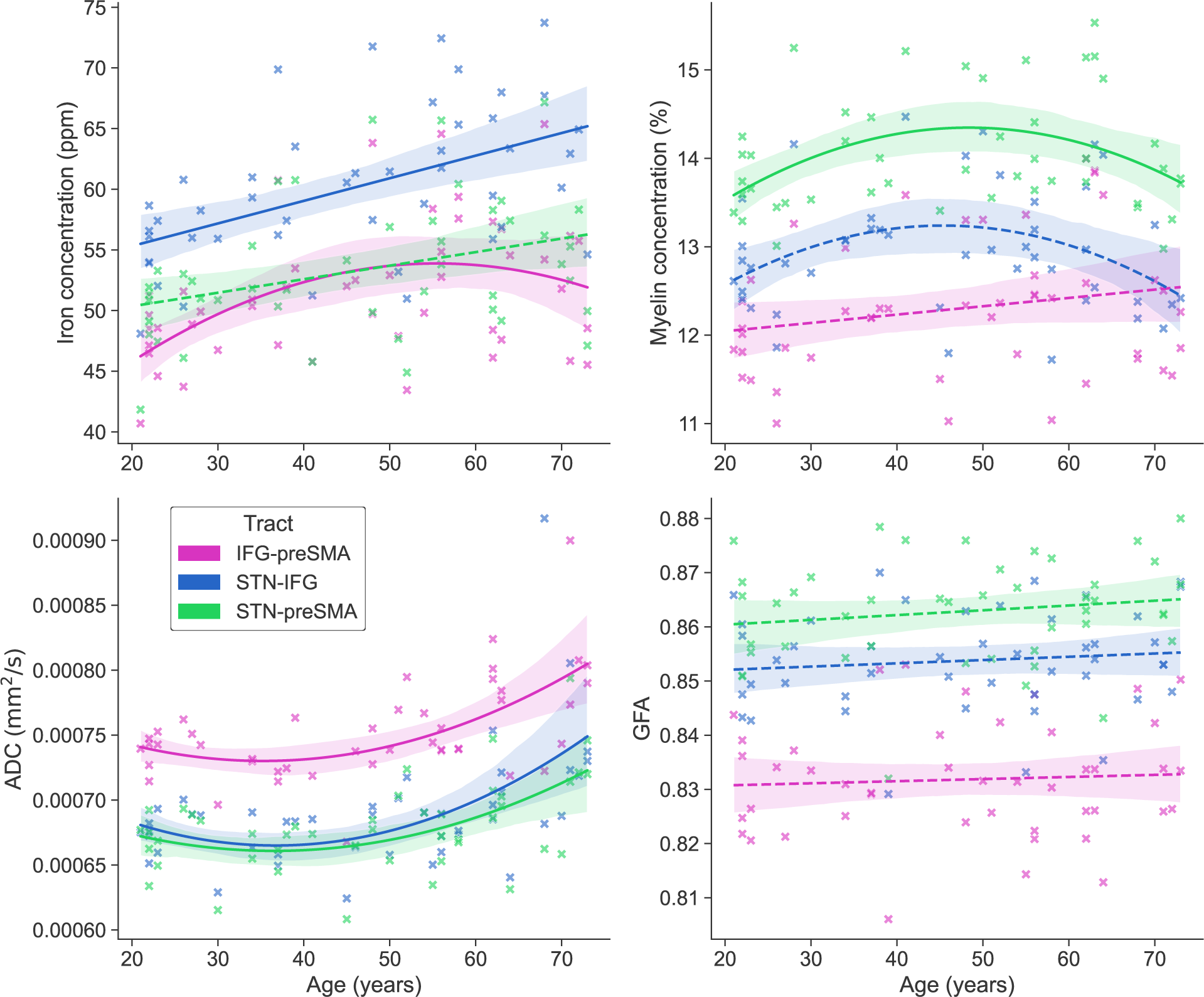
Age-related changes in the IFG-preSMA, STN-IFG, and STN-preSMA tracts in iron (top left), myelin (top right), apparent diffusion coefficient (ADC; bottom left), and generalised fractional anisotropy (GFA; bottom right). Note that some models had a relatively low *R*^2^; instances where *R*^2^ < 0.15 are denoted with a dashed line. See Table 2 for details. Error bars depict confidence interval of the regression line.

#### 3.3.2 Mediation of iron and myelin

To further investigate the relationship between these variables, we assessed the influence of iron and myelin on the relationship between age and the diffusion measures via a mediation analysis, for each tract of interest. The mediation results are given briefly here, but estimates and significance tests for direct, indirect, and total effects for each tract can be found in the Supplementary Materials.

#### ADC

The mediation analyses found that iron and myelin were almost never significant mediators of the relationship between age and ADC. The indirect relationship was only significant for the IFG-preSMA for iron (*p* = .030), but the estimates for the direct effect were much larger (indirect effect: -0.042, *p* = .030, ADE: 0.322, *p < .*001), indicating that the relationship between ADC and age is quite strong without iron.

#### GFA

There were no mediation effects for iron or myelin for GFA for any of the tracts of interest (all *p > .*550).

## 4 Discussion

The STN, IFG, and preSMA play distinct roles in action cancellation. Current models of response inhibition propose that these regions, among others, operate as a network, coordinating activity in order to successfully cancel or modify volitional movement. This inhibitory ability is compromised in older populations, with older adults being slower to cancel their movements. In the current study, we quantified anatomical changes that occur in the STN, IFG, and preSMA and their connecting pathways by combining high-resolution DWI and ultra-high field qMRI in the same participants, quantifying changes using polynomial regression. To our knowledge, this is this first study to examine age-related biophysical and structural changes in the stopping network.

Broadly, we identified substantial age-related anatomical change at the network level, i.e., in the tracts that connect these regions, with less evidence for localised change, i.e., anatomical change in the regions themselves. Taken together, the results suggest that the behavioural changes in response inhibition efficacy and altered neural recruitment patterns may be the product of network-level breakdown, or connectivity alterations.

### 4.1 Minimal localised changes (ROIs)

We found the age-related changes in iron or myelin to be mostly linear for the regions of interest (IFG, preSMA, STN), however it should be noted that these models had generally low *R*^2^ values, indicating that the age-related differences in these values is low relative to other sources of inter-individual variability.

Age-related changes in iron concentration have previously been observed in the STN, although not always (for a review, see Madden and Merenstein, 2023). It should be noted that estimation methods for iron vary. Here, we used a combination of QSM and R2* to estimate iron; previous work has often used QSM or R2* individually as proxies for iron (see e.g, Betts et al., 2016; Burgetova et al., 2021; Keuken et al., 2017). The sample size of the current study was limited due to the availability of DWI data and we have previously observed age-related iron accumulation in the STN in larger samples (Miletić et al., 2022). Further, much previous work has focused on clinical samples and particularly, identified iron accumulation patterns in clinical samples late in the disease course. Iron accumulation in the STN has been correlated with severe motor symptoms in late-stage Parkinson’s disease (PD) and has been hypothesised to be an important predictor of therapeutic and surgical outcomes (Brown et al., 2022; Huang et al., 2020). However, in early disease stages this association is less clear (Shin et al., 2018) and in healthy ageing contexts, the age-related changes can be inconsistent (Madden & Merenstein, 2023). Collectively, these findings suggest that local iron change may exhibit subtle variations, and the relationship between local anatomical changes and functional or behavioural change may be more difficult to describe, especially compared to broader network changes (see below, 4.2).

Turning to the cortex, to our knowledge there are no studies that have specifically examined age-related changes in iron in the cortex in non-clinical populations. However, Burgetova and colleagues (2021) did identify age-related changes in magnetic susceptibility in broader premotor and frontal regions, with these susceptibility changes assumed to be influenced by iron concentration. Varied broader cortical changes in some qMRI metrics in older adult populations have also been observed (Gracien et al., 2017; Seiler et al., 2020), including in prefrontal areas (Gozdas et al., 2021), but again, these are not particularly region-specific. Although there are some limitations to our approach (see 4.5), this highlights the difficulty in capturing specific anatomical change even in relatively large cortical areas.

### 4.2 Age-related changes in the stopping network

We observed quadratic and linear increases in iron in the tracts of interest (IFG-preSMA, STN-IFG, and STN-preSMA), quadratic and linear changes in myelin, and and quadratic increases in ADC. The GFA models for all three tracts of interest had an extremely small *R*^2^, indicating minimal age-related change in this metric.

#### 4.2.1 Iron

Iron is increasingly recognised as a significant factor in both diseased and healthy ageing contexts. While it is an essential component of neural functioning, iron is also an active oxidiser. Excessive iron accumulation is thought to be a factor in tissue degeneration, interrupting normal cellular functioning and causing degradation of cellular components such as mitochondria, lipids, and proteins (Daugherty & Raz, 2015; Hagemeier et al., 2012). Its accumulation has also been proposed to impact other neurophysiological measures, such as neural oscillatory patterns in the beta frequency band (Lin et al., 2023). Lin and colleagues propose that in a PD context, observed alterations in beta oscillatory activity are caused by excessive iron accumulation. This accumulation causes dopaminergic cell death in critical motor-related regions, affecting subcortical activity mediated by acetylcholine and *γ*-aminobutyric acid (GABA) and causing alterations in the intricate inhibitory circuitry of the subcortex. This ultimately leads to hyperactivation of the STN and the observed increase in beta oscillations. Beta oscillations have also been tightly coupled with stopping behaviour (Wessel, 2020), with beta band coherence between frontal and motor regions (including the preSMA and IFG) correlated with stopping efficacy (Ding et al., 2023). GABAergic activity has also been linked to response inhibition in both younger and older adults, and older adults tend to show a reduction of GABAergic levels and activity (Faßbender et al., 2021; Hermans et al., 2019; Quetscher et al., 2015; Verstraelen et al., 2021). Here, we have shown patterns of iron accumulation in the stopping network across the adult lifespan. Tentatively, this accumulation may be a critical component of the neural breakdown that occurs in this network in older age, which ultimately leads to the aforementioned observed neural and behavioural changes, including alterations to beta oscillatory activity and GABergic activity, and SSRT slowing.

#### 4.2.2 Myelin

Like iron, myelin is essential for healthy brain function, speeding signal conduction and facilitating longer-range signal transmission (Radtke et al., 2007; Williamson & Lyons, 2018; Yeatman et al., 2014). Changes in myelin are associated with typical healthy ageing, with a loss of myelin sheaths and myelinated nerve fibres affecting the functioning of neural circuits and contributing to normal cognitive decline (Peters, 2009; X. Zhang et al., 2021). Altered demyelination patterns have also been observed in a number of neurodegenerative conditions such as AD and multiple sclerosis (Bartzokis, 2011; Grussu et al., 2017; Radtke et al., 2007). These demyelination patterns have been proposed to be a critical element of the neural breakdown that occurs in AD (incidentally, stimulated by iron accumulation; see Bartzokis, 2011; Bartzokis et al., 2007; Khattar et al., 2021). Previous work has identified broad quadratic decreases in myelin with older age, similar to those observed in the current study (Buyanova & Arsalidou, 2021; Khattar et al., 2021; Miletić et al., 2022). Note that just like in the case of iron, different MRI metrics have been used as proxies for myelin. For example, Yeatman and colleagues (2014) used R1, Khattar and colleagues (2021) used myelin water fraction, and Grydeland and colleagues (2019) used the T1/T2 ratio. Whilst these estimates have been associated with myelination, they have also been demonstrated to be impacted by other factors (e.g., the T1/T2 ratio is more strongly associated with axon density than myelin density; Grydeland et al., 2019). Comparison of findings should, therefore, be made with due caution. Here (using a combination of R1 and R2*), we observed some patterns of demyelination in cortical-subcortical tracts, but particularly in the STN-preSMA tract.

Although the IFG has historically been posited as being the site for action cancellation (Aron et al., 2014; Suda et al., 2020), the role of the IFG in attentional monitoring has also been emphasised (Chatham et al., 2012; Munakata et al., 2011; Sharp et al., 2010). Under the recently-proposed Pause-then-Cancel (PTC) model, the role of these two regions has been further demarcated, with the IFG proposed to play a generic attentional-monitoring role (the “pause” process) and the preSMA being the cortical region involved in action cancellation, realised via the subcortex (Diesburg & Wessel, 2021). Importantly, it is action cancellation specifically that is compromised in older adult populations; more proactive or attentional elements of stopping performance are relatively well-preserved in older age (Bloemendaal et al., 2016; Hermans et al., 2019; Kleerekooper et al., 2016). Speculatively, the decrease in myelination observed in this STN-preSMA tract may represent some neuroanatomical changes that underpin these behavioural changes.

#### 4.2.3 ADC and FA

We observed consistent quadratic increases in the stopping network in ADC, but not in GFA. With respect to response inhibition networks, work to date has examined the relationship between response inhibition networks and FA, rather than GFA, but these findings have been highly variable. For example, Madsen and colleagues (2020) found FA correlated with SSRT in the preSMA only (i.e., not in the IFG or in any subcortical regions), whereas Coxon and colleagues (2012) found this association in the IFG. Boen and colleagues (2022) found an association between stopping accuracy and FA in an IFG subcomponent^1^-preSMA tract only (other regions of interest included the caudate, putamen, insula, and STN). These estimations of FA have largely been done using a tensor estimation method with relatively simple protocols, which has a number of limitations. An extensive comparison of estimation techniques is beyond the scope of this paper, but briefly, these methods assume that diffusivity in each voxel can be described in a single direction. This creates problems in instances of crossing fibres (which may occur in up to 90% of all voxels: Jeurissen et al., 2013; Schilling et al., 2021) and favours the identification of major connectivity pathways, limiting reconstruction of more minor pathways (Dell’Acqua & Tournier, 2019; Volz et al., 2018). Conversely, GFA is useful for estimating diffusivity in more complex tissue environments, i.e., in regions of high curvature or where there are many crossing fibres (Glenn et al., 2015; Sun et al., 2023). Although like FA, GFA does correlate negatively with age (Porcu et al., 2021), previous work has found these changes to be highly specific from region to region, meaning generalised statements about age-associated changes should be considered with caution (Teipel et al., 2014). Moreover, changes to white matter structure may have varying impact on these metrics, particularly FA. They quantify relative diffusivity, meaning that if degradation to white matter causes uniform changes to diffusivity, FA can be largely unchanged despite gross changes in the white matter (Figley et al., 2022). Additionally, in tensor estimation approaches, only the diffusivity in the major direction is quantified. If there are increases in white matter density in one of the non-major directions, this will lower the anisotropy in that voxel, which will paradoxically lower FA, when there has actually been an *increase* in white matter density (Figley et al., 2022). This indicates that while FA is sensitive to anatomical changes, its interpretation in terms of specific causes remains challenging (Jones & Cercignani, 2010). Instead, metrics which assess net diffusion, i.e., not diffusion in a particular direction, may provide more sensitive estimations.

Converse to the GFA results, for ADC (a measure of net diffusion: Helenius et al., 2002), we found consistent age-related increases in all tracts of interest. In young adults, mean diffusivity has been found to positively correlate with SSRT in STN-IFG and STN-preSMA tracts (Rae et al., 2015), indicating that white matter integrity is intrinsic to action cancellation efficacy. Here, we observed an age-associated increase in ADC, which may be reflective of alterations in the underlying anatomy (importantly, these changes probably more reliably reflect anatomical degradation, as ADC has been demonstrated to be more sensitive to underlying anatomical changes than FA: Scheurer et al., 2011).

### 4.3 Models of ageing

Theoretical frameworks of ageing such as STAC and CRUNCH postulate that functional recruitment changes observed in older adult populations are preceded by measurable anatomical changes. For example, the STAC model hypothesises that functional recruitment changes are preceded by anatomical change including brain volume reduction, connectivity/white matter changes, cortical thinning, and dopamine depletion (Park & Reuter-Lorenz, 2009; Reuter-Lorenz & Park, 2014). These changes results in broader recruitment patterns, which compensate for the structural changes by recruiting a wider range of neural regions than was previously required. Here, we have shown, for the first time, evidence of complex anatomical changes specific to the stopping network, and suggest that these may precede or underpin the well-documented functional and behavioural changes in response inhibition in older adult populations.

Crucially, the majority of these anatomical changes were observable in the tracts that connect the canonical stopping regions, and not the regions themselves, despite these regions showing blood-oxygenation level dependent (BOLD) activity changes in older adult populations (Bloemendaal et al., 2016; Coxon et al., 2016). We suggest that it may be the connectivity changes that most crucially underpin these observed functional changes or at least, are the most broadly measurable. This is a particularly salient point given the requirement for fast and efficacious transmission of information in subcortical-cortical pathways under response inhibition frameworks (Aron, 2007; Diesburg & Wessel, 2021; Narayanan et al., 2020), which may deteriorate in older age.

It should also be noted that variation in localised anatomical changes (i.e., in individual regions) are thought to be the cause of the substantial individual differences in age-related cognitive change (Kang et al., 2022). Likewise, in the context of response inhibition, a range of anatomical measures have been found to modulate individual performance (see e.g., Boy et al., 2010; Chowdhury et al., 2019; Coxon et al., 2016; Forstmann et al., 2012; King et al., 2012). More localised anatomical changes, e.g., changes to the stopping regions themselves, were not particularly evident in the current study, but may be better elucidated with the use of behavioural data, where anatomical changes could be linked with individual cognitive or behavioural measures.

### 4.4 Myelin and iron do not mediate diffusion indices

DWI is a way of estimating the structure of the underlying white matter, based on the assumption that the movement in the white matter is more constrained than in the grey matter or cerebrospinal fluid. It is assumed that demyelination affects these measures, resulting in increases in diffusivity (i.e., the movement of water is less constrained). There are few studies that have correlated diffusion measures with non-DWI approximations of myelin or *ex vivo* measures, particularly outside of a clinical context, meaning the extent to which these metrics do truly index myelin remains unclear (Arshad et al., 2016; Henriques et al., 2023; Lazari & Lipp, 2021).

Here, we first identified relationships between age and the diffusion metrics, and then used a mediation approach to assess if iron and myelin were mediators of these relationships. We largely found no mediation effects (except for one instance, but the direct effect was still larger than the indirect effect, i.e., the mediation effect was comparatively small). Given we are only examining a small number of tracts, these results should not be taken as representative of the whole brain. However, it does demonstrate that, particularly in an ageing context, diffusion metrics may not directly reflect myelination. This is perhaps unsurprising given the huge number of anatomical changes that take place in older age; while demyelination and iron accumulation are certainly a part of these, there are also broader anatomical changes including a loss of dendritic spines, decreases in brain volume, and an alterations of astroctytes and other microglia, (Palmer & Ousman, 2018; Yeatman et al., 2014), which more broadly affects the central nervous system and will also impact diffusivity. Thus, whilst diffusion metrics can be a highly useful metric for studying anatomical change, direct interpretations of the specific changes that they index should be made with caution.

### 4.5 Limitations

The present study has several limitations. Firstly, the data in the present study are purely anatomical, i.e., we have no behavioural data for the participants. Behavioural data was collected for this sample, but due to COVID-19 restrictions, was gathered online, resulting in a small sample size and insufficient data quality for further analyses (the MRI data had been acquired prior to this time). Instead, we relied brain-behaviour relations reported in the literature to inform our interpretations. Future work should correlate these metrics (behavioural, qMRI, and DWI) in the same sample to strengthen the findings, particularly given the emphasis on individual differences known to affect ageing and stop signal performance (Chowdhury et al., 2019; Forstmann et al., 2012; Isherwood, Bazin, et al., 2023).

Secondly, the iron and myelin metrics used in this sample are approximations. There is no direct way to measure iron and myelin content in vivo but as discussed above, there are a number of different ways to approximate these metrics. In the current study, we used simplified biophysical linear models that estimated iron and myelin concentrations using a combination of qMRI contrast values. These models were selected from eight model candidates, with all possible combinations of R1, R2*, and QSM parameters. Their predictability was verified using literature-based concentrations (for details, see Miletić et al., 2022), but are proxies nonetheless.

Thirdly, imaging acquisition and assessment of brain images is complicated in aged samples. Anatomical changes that occur in older age are highly variable and can cause difficulties in applying standard pipelines or e.g., decrease parcellation accuracy (Alkemade et al., 2022; Bazin et al., 2020). Further, MRI values can be independently affected by age-related alterations, which confound data and make it difficult to isolate specific anatomical changes. For example, older adults tend to have white matter hyperintensities, which in turn affects diffusion metrics. These hyperintensities can be identified using fluid attenuated inversion recovery (FLAIR) scans (Tubi et al., 2020), but these were not part of the acquisition sequence in the present study.

Finally, we chose to focus on three regions of interest in the present study. While these regions are well-documented as being intrinsic to action cancellation (see e.g., Aron et al., 2014; Borgomaneri et al., 2020; Chen et al., 2020; Lee et al., 2016; Lofredi et al., 2021), there are many additional regions that have been associated with response inhibition, particularly in the subcortex. Theoretical models of stopping such as the PTC model postulate that response inhibition involves a complex interplay of cortical and subcortical regions including the substantia nigra, globus pallidus, and striatum (Diesburg & Wessel, 2021; R. Schmidt & Berke, 2017; R. Schmidt et al., 2013), with experimental evidence also identifying the ventral tegmental area and thalamus (Isherwood, Kemp, et al., 2023; Tennyson et al., 2018). Due to difficulties imaging the subcortex (de Hollander et al., 2017; Miletić et al., 2020), the specific role of these regions remains undetermined; more research is required to disentangle their specific role in action cancellation. Future work could include a broader range of anatomical regions in order to more fully encompass the response inhibition-associated regions.

## 5 Conclusion

A large body of imaging and brain stimulation research has identified the IFG, preSMA, and STN as being intrinsic to action cancellation. These regions, among others, form a stopping ‘network’, and are thought to coordinate to realise action selection and cancellation through a series of subcortical-cortical pathways. The connectivity of these regions is known to be an important factor in action cancellation and stopping, with improved connectivity of these regions associated with increased performance in the stop-signal task.

Here, we quantified the anatomical changes that occur in these regions across the lifespan and in the white matter pathways that connect them, combining high-resolution DWI and ultra-high field qMRI in the same sample. We found substantial changes in iron concentration in these tracts, increases in ADC, and some evidence for demyelination. Conversely, we found very little evidence for age-related anatomical changes in the regions themselves. We propose that some of the functional changes observed in these regions in older adult populations (e.g., increased BOLD recruitment) are a reflection of alterations to the stopping network itself, i.e., to connectivity changes, rather than to localised regional change.

## Data and code availability statement

The preprocessed DWI and qMRI datasets have been previously published and are publicly available (Alkemade et al., 2020; Keuken et al., 2022). All code used to estimate the tractography, run analyses, and generate figures can be found at https://osf.io/hnz79/.

## CRediT author contributions

**Sarah A Kemp:** Conceptualisation, Methodology, Software, Validation, Formal analysis, Data curation, Writing – Original draft, Writing – Review & editing, Visualisation. **Pierre-Louis Bazin:** Conceptualisation, Methodology, Software, Validation, Investigation, Data curation, Writing – Review & editing, Supervision, Project administration. **Steven Miletić:** Investigation, Data curation, Software, Validation, Writing – Review & editing. **Russell J Boag:** Validation, Writing – Review & editing. **Max C Keuken:** Conceptualisation, Software, Investigation, Data curation, Supervision. **Mark R Hinder:** Supervision. **Birte U Forstmann:** Conceptualisation, Resources, Writing – Review & editing, Supervision, Project administration, Funding acquisition.

## Acknowledgements

This research was financially supported by STW/NWO (#14017; BUF), ERC Consolidator (BUF), ERC PoC (BUF), NWO Vici (016.Vici.185.052; BUF), Stichting Ammodo (2019; BUF), and a University of Tasmania Graduate Research Scholarship (SAK). The authors thank Niek Stevenson for helpful discussions regarding data analysis on this manuscript.

## Footnotes

1 IFG dorsal pars opercularis

## References

Alkemade, A., Mulder, M. J., Groot, J. M., Isaacs, B. R., van Berendonk, N., Lute, N., Isherwood, S. J., Bazin, P.-L., & Forstmann, B. U. (2020). The Amsterdam Ultra-high field adult lifespan database (AHEAD): A freely available multimodal 7 Tesla submillimeter magnetic resonance imaging database. NeuroImage, 221, 117200. 10.1016/ j.neuroimage.2020.117200

Alkemade, A., Mulder, M. J., Trutti, A. C., & Forstmann, B. U. (2022). Manual delineation approaches for direct imaging of the subcortex. Brain Structure and Function, 227(1), 219–297. 10.1007/s00429-021-02400-x

Aron, A. R. (2007). The neural basis of inhibition in cognitive control. The Neuroscientist, 13(3). 10.1177/1073858407299288

Aron, A. R., Behrens, T. E., Smith, S., Frank, M. J., & Poldrack, R. A. (2007). Triangulating a cognitive control network using diffusion-weighted magnetic resonance imaging (MRI) and functional MRI. Journal of Neuroscience, 27(14), 3743–3752. 10.1523/ JNEUROSCI.0519-07.2007

Aron, A. R., & Poldrack, R. A. (2006). Cortical and subcortical contributions to stop signal response inhibition: Role of the subthalamic nucleus. Journal of Neuroscience, 26(9), 2424– 2433. 10.1523/JNEUROSCI.4682-05.2006

Aron, A. R., Robbins, T. W., & Poldrack, R. A. (2004). Inhibition and the right inferior frontal cortex. Trends in Cognitive Sciences, 8(4), 170–177. 10.1016/j.tics.2004.02.010

Aron, A. R., Robbins, T. W., & Poldrack, R. A. (2014). Inhibition and the right inferior frontal cortex: One decade on. Trends in Cognitive Sciences, 18(4), 177–185. 10.1016/j.tics.2013.12.003

Arshad, M., Stanley, J. A., & Raz, N. (2016). Adult age differences in subcortical myelin content are consistent with protracted myelination and unrelated to diffusion tensor imaging indices. NeuroImage, 143, 26–39. 10.1016/j.neuroimage.2016.08.047

Bartzokis, G. (2011). Alzheimer’s disease as homeostatic responses to age-related myelin break-down. Neurobiology of aging, 32(8), 1341–1371. 10.1016/j.neurobiolaging.2009.08.007

Bartzokis, G., Tishler, T. A., Lu, P. H., Villablanca, P., Altshuler, L. L., Carter, M., Huang, D., Edwards, N., & Mintz, J. (2007). Brain ferritin iron may influence age- and gender-related risks of neurodegeneration. Neurobiology of Aging, 28(3), 414–423. 10.1016/j.neurobiolaging.2006.02.005

Bazin, P.-L., Alkemade, A., Mulder, M. J., Henry, A. G., & Forstmann, B. U. (2020). Multi-contrast anatomical subcortical structures parcellation. eLife, 9, e59430. 10.7554/ eLife.59430

Bazin, P.-L., Alkemade, A., van der Zwaag, W., Caan, M., Mulder, M., & Forstmann, B. U. (2019). Denoising high-field multi-dimensional MRI with local complex PCA. Frontiers in Neuroscience, 13, 1066. 10.3389/fnins.2019.01066

Beck, D., de Lange, A.-M. G., Maximov, I. I., Richard, G., Andreassen, O. A., Nordvik, J. E., & Westlye, L. T. (2021). White matter microstructure across the adult lifespan: A mixed longitudinal and cross-sectional study using advanced diffusion models and brain-age prediction. NeuroImage, 224, 117441. 10.1016/j.neuroimage.2020.117441

Bedard, A.-C., Nichols, S., Barbosa, J., Schachar, R., Logan, G., & Tannock, R. (2002). The development of selective inhibitory control across the life span. Developmental Neuropsychology, 21(1), 93–111. 10.1207/S15326942DN2101_5

Betts, M. J., Acosta-Cabronero, J., Cardenas-Blanco, A., Nestor, P. J., & Düzel, E. (2016). High-resolution characterisation of the aging brain using simultaneous quantitative susceptibility mapping (QSM) and R2* measurements at 7T. NeuroImage, 138, 43–63. 10.1016/j.neuroimage.2016.05.024

Bingham, C. S., Petersen, M. V., Parent, M., & McIntyre, C. C. (2023). Evolving characterization of the human hyperdirect pathway. Brain Structure and Function, 228(2), 353–365. 10.1007/s00429-023-02610-5

Bissett, P. G., & Logan, G. D. (2011). Balancing cognitive demands: Control adjustments in the stop-signal paradigm. *Journal of Experimental Psychology: Learning*, Memory, and Cognition, 37(2), 392–404. 10.1037/a0021800

Bloemendaal, M., Zandbelt, B., Wegman, J., van de Rest, O., Cools, R., & Aarts, E. (2016). Contrasting neural effects of aging on proactive and reactive response inhibition. Neurobiology of Aging, 46, 96–106. 10.1016/j.neurobiolaging.2016.06.007

Boen, R., Raud, L., & Huster, R. J. (2022). Inhibitory control and the structural parcelation of the right inferior frontal gyrus. Frontiers in Human Neuroscience, 16, 787079. 10.3389/fnhum.2022.787079

Borgomaneri, S., Serio, G., & Battaglia, S. (2020). Please, don’t do it! Fifteen years of progress of non-invasive brain stimulation in action inhibition. Cortex, 132, 404–422. 10.1016/j.cortex.2020.09.002

Boy, F., Evans, C. J., Edden, R. A., Singh, K. D., Husain, M., & Sumner, P. (2010). Individual differences in subconscious motor control predicted by GABA concentration in SMA. Current Biology, 20(19), 1779–1785. 10.1016/j.cub.2010.09.003

Brown, G., Du, G., Farace, E., Lewis, M., Eslinger, P., McInerney, J., Kong, L., Li, R., Huang, X., & De Jesus, S. (2022). Subcortical iron accumulation pattern may predict neuropsychological outcomes after subthalamic nucleus deep brain stimulation: A pilot study. Journal of Parkinson’s Disease, 12(3), 851–863. 10.3233/JPD-212833

Burgetova, R., Dusek, P., Burgetova, A., Pudlac, A., Vaneckova, M., Horakova, D., Krasensky, J., Varga, Z., & Lambert, L. (2021). Age-related magnetic susceptibility changes in deep grey matter and cerebral cortex of normal young and middle-aged adults depicted by whole brain analysis. Quantitative Imaging in Medicine and Surgery, 11(9), 3906–3919. 10.21037/qims-21-87

Buyanova, I. S., & Arsalidou, M. (2021). Cerebral white matter myelination and relations to age, gender, and cognition: A selective review. Frontiers in Human Neuroscience, 15. https ://doi.org/10.3389/fnhum.2021.662031

Caan, M. W. A., Bazin, P.-L., Marques, J. P., de Hollander, G., Dumoulin, S. O., & van der Zwaag, W. (2019). MP2RAGEME: T1, T2*, and QSM mapping in one sequence at 7 tesla. Human Brain Mapping, *40*(6), 1786–1798. 10.1002/hbm.24490

Cavanagh, J. F., Sanguinetti, J. L., Allen, J. J. B., Sherman, S. J., & Frank, M. J. (2014). The subthalamic nucleus contributes to post-error slowing. Journal of Cognitive Neuroscience, 26(11), 2637–2644. 10.1162/jocn_a_00659

Chatham, C. H., Claus, E. D., Kim, A., Curran, T., Banich, M. T., & Munakata, Y. (2012). Cognitive control reflects context monitoring, not motoric stopping, in response inhibition. PLOS ONE, 7(2), e31546. 10.1371/journal.pone.0031546

Chen, W., de Hemptinne, C., Miller, A. M., Leibbrand, M., Little, S. J., Lim, D. A., Larson, P. S., & Starr, P. A. (2020). Prefrontal-subthalamic hyperdirect pathway modulates movement inhibition in humans. Neuron, 106(4), 579–588.e3. 10.1016/j.neuron.2020.02.012

Chowdhury, N. S., Livesey, E. J., & Harris, J. A. (2019). Individual differences in intracortical inhibition during behavioural inhibition. Neuropsychologia, 124, 55–65. 10.1016/j.neuropsychologia.2019.01.008

Coudé, D., Parent, A., & Parent, M. (2018). Single-axon tracing of the corticosubthalamic hyperdirect pathway in primates. Brain Structure and Function, 223(9), 3959–3973. 10.1007/s00429-018-1726-x

Coxon, J. P., Goble, D. J., Leunissen, I., Van Impe, A., Wenderoth, N., & Swinnen, S. P. (2016). Functional brain activation associated with inhibitory control deficits in older adults. Cerebral Cortex, 26(1), 12–22. 10.1093/cercor/bhu165

Coxon, J. P., Van Impe, A., Wenderoth, N., & Swinnen, S. P. (2012). Aging and inhibitory control of action: Cortico-subthalamic connection strength predicts stopping performance. The Journal of Neuroscience, 32(24), 8401–8412. 10.1523/JNEUROSCI.6360-11.2012

Daugherty, A. M., & Raz, N. (2015). Appraising the role of iron in brain aging and cognition: Promises and limitations of MRI methods. Neuropsychology Review, 25(3), 272–287. 10.1007/s11065-015-9292-y

de Hollander, G., Keuken, M., van der Zwaag, W., Forstmann, B., & Trampel, R. (2017). Comparing functional MRI protocols for small, iron-rich basal ganglia nuclei such as the subthalamic nucleus at 7 T and 3 T. Human Brain Mapping, 38(6), 3226–3248. 10.1002/hbm.23586

Dell’Acqua, F., & Tournier, J.-D. (2019). Modelling white matter with spherical deconvolution: How and why? NMR in Biomedicine, 32(4), e3945. 10.1002/nbm.3945

Diesburg, D. A., & Wessel, J. (2021). The Pause-then-Cancel model of human action stopping: Theoretical considerations and empirical evidence. Neuroscience & Biobehavioral Reviews, 129, 17–34. 10.31234/osf.io/vp6es

Ding, Q., Lin, T., Cai, G., Ou, Z., Yao, S., Zhu, H., & Lan, Y. (2023). Individual differences in beta-band oscillations predict motor-inhibitory control. Frontiers in Neuroscience, 17. 10.3389/fnins.2023.1131862

Eagle, D. M., Baunez, C., Hutcheson, D. M., Lehmann, O., Shah, A. P., & Robbins, T. W. (2008). Stop-signal reaction-time task performance: Role of prefrontal cortex and subthalamic nucleus. Cerebral Cortex, 18(1), 178–188. 10.1093/cercor/bhm044

Faßbender, K., Bey, K., Lippold, J. V., Aslan, B., Hurlemann, R., & Ettinger, U. (2021). GABAergic modulation of performance in response inhibition and interference control tasks. Journal of Psychopharmacology, 35(12), 1496–1509. 10.1177/02698811211032440

Figley, C. R., Uddin, M. N., Wong, K., Kornelsen, J., Puig, J., & Figley, T. D. (2022). Potential pitfalls of using fractional anisotropy, axial diffusivity, and radial diffusivity as biomarkers of cerebral white matter microstructure. Frontiers in Neuroscience, 15. 10.3389/fnins.2021.799576

Forstmann, B. U., Keuken, M. C., Jahfari, S., Bazin, P.-L., Neumann, J., Schäfer, A., Anwander, A., & Turner, R. (2012). Cortico-subthalamic white matter tract strength predicts interindividual efficacy in stopping a motor response. NeuroImage, 60(1), 370–375. 10.1016/j.neuroimage.2011.12.044

Frank, M. J. (2006). Hold your horses: A dynamic computational role for the subthalamic nucleus in decision making. Neural Networks: The Official Journal of the International Neural Network Society, 19(8), 1120–1136. 10.1016/j.neunet.2006.03.006

Geerligs, L., Renken, R. J., Saliasi, E., Maurits, N. M., & Lorist, M. M. (2015). A brain-wide study of age-related changes in functional connectivity. Cerebral Cortex, 25(7), 1987– 1999. 10.1093/cercor/bhu012

Glenn, G. R., Helpern, J. A., Tabesh, A., & Jensen, J. H. (2015). Quantitative Assessment of Diffusional Kurtosis Anisotropy. NMR in Biomedicine, 28(4), 448–459. 10.1002/ nbm.3271

Gozdas, E., Fingerhut, H., Wu, H., Bruno, J. L., Dacorro, L., Jo, B., O’Hara, R., Reiss, A. L., & Hosseini, S. M. H. (2021). Quantitative measurement of macromolecular tissue properties in white and gray matter in healthy aging and amnestic MCI. NeuroImage, 237, 118161. 10.1016/j.neuroimage.2021.118161

Gracien, R.-M., Nürnberger, L., Hok, P., Hof, S.-M., Reitz, S. C., Rüb, U., Steinmetz, H., Hilker-Roggendorf, R., Klein, J. C., Deichmann, R., & Baudrexel, S. (2017). Evaluation of brain ageing: A quantitative longitudinal MRI study over 7 years. European Radiology, 27(4), 1568–1576. 10.1007/s00330-016-4485-1

Graybiel, A. M. (2000). The basal ganglia. Current Biology, 10(14), R509–R511. 10.1016/S0960-9822(00)00593-5

Grier, M. D. (2020). Estimating brain connectivity with diffusion-weighted MRI: Promise and peril. Biological Psychiatry: Cognitive Neuroscience and Neuroimaging, 5(9), 846–854. 10.1016/j.bpsc.2020.04.009

Grussu, F., Schneider, T., Tur, C., Yates, R. L., Tachrount, M., Ianuş, A., Yiannakas, M. C., Newcombe, J., Zhang, H., Alexander, D. C., DeLuca, G. C., & Gandini Wheeler-Kingshott, C. A. M. (2017). Neurite dispersion: A new marker of multiple sclerosis spinal cord pathology? Annals of Clinical and Translational Neurology, 4(9), 663–679. 10.1002/acn3.445

Grydeland, H., Vértes, P. E., Váša, F., Romero-Garcia, R., Whitaker, K., Alexander-Bloch, A. F., Bjørnerud, A., Patel, A. X., Sederevičius, D., Tamnes, C. K., Westlye, L. T., White, S. R., Walhovd, K. B., Fjell, A. M., & Bullmore, E. T. (2019). Waves of maturation and senescence in micro-structural MRI markers of human cortical myelination over the lifespan. Cerebral Cortex, 29(3), 1369–1381. 10.1093/cercor/bhy330

Hagemeier, J., Geurts, J. J., & Zivadinov, R. (2012). Brain iron accumulation in aging and neurodegenerative disorders. Expert Review of Neurotherapeutics, 12(12), 1467–1480. 10.1586/ern.12.128

Hagmann, P., Cammoun, L., Gigandet, X., Meuli, R., Honey, C. J., Wedeen, V. J., & Sporns, O. (2008). Mapping the structural core of human cerebral cortex. PLOS Biology, 6(7), e159. 10.1371/journal.pbio.0060159

Healey, R., Goldsworthy, M., Salomoni, S., Weber, S., Kemp, S., Hinder, M. R., & St George, R. J. (2024). Impaired motor inhibition during perceptual inhibition in older, but not younger adults: A psychophysiological study. Scientific Reports, 14(1), 2023. 10.1038/s41598-024-52269-z

Helenius, J., Soinne, L., Perkiö, J., Salonen, O., Kangasmäki, A., Kaste, M., Carano, R. A. D., Aronen, H. J., & Tatlisumak, T. (2002). Diffusion-weighted MR imaging in normal human brains in various age groups. AJNR: American Journal of Neuroradiology, 23(2), 194. https://www.ncbi.nlm.nih.gov/pmc/articles/PMC7975251/

Henriques, R. N., Henson, R., Cam-CAN, & Correia, M. M. (2023). Unique information from common diffusion MRI models about white-matter differences across the human adult lifespan. Imaging Neuroscience, 1, 1–25. 10.1162/imag_a_00051

Hermans, L., Maes, C., Pauwels, L., Cuypers, K., Heise, K.-F., Swinnen, S. P., & Leunissen, I. (2019). Age-related alterations in the modulation of intracortical inhibition during stopping of actions. Aging, 11(2), 371–385. 10.18632/aging.101741

Hsieh, S., & Lin, Y.-C. (2017). Stopping ability in younger and older adults: Behavioral and event-related potential. *Cognitive, Affective*, & Behavioral Neuroscience, 17(2), 348–363. 10.3758/s13415-016-0483-7

Hu, S., Job, M., Jenks, S., Chao, H., & Li, C.-S. (2019). Imaging the effects of age on proactive control in healthy adults. Brain Imaging and Behavior, 13(6), 1526–1537. 10.1007/s11682-019-00103-w

Hu, S., Ide, J. S., Chao, H. H., Castagna, B., Fischer, K. A., Zhang, S., & Li, C.-s. R. (2018). Structural and functional cerebral bases of diminished inhibitory control during healthy aging. Human Brain Mapping, 39(12), 5085–5096. 10.1002/hbm.24347

Huang, W., Ogbuji, R., Zhou, L., Guo, L., Wang, Y., & Kopell, B. H. (2020). Motoric impairment versus iron deposition gradient in the subthalamic nucleus in Parkinson’s disease [Publisher: American Association of Neurological Surgeons Section: Journal of Neurosurgery]. Journal of Neurosurgery, 135(1), 284–290. 10.3171/2020.5.JNS201163

Huntenburg, J. M., Steele, C. J., & Bazin, P.-L. (2018). Nighres: Processing tools for high-resolution neuroimaging. GigaScience, 7(7), giy082. 10.1093/gigascience/giy082

Imai, K., Keele, L., & Tingley, D. (2010). A general approach to causal mediation analysis. Psychological Methods, 15(4), 309–334. 10.1037/a0020761

Isherwood, S. J. S., Bazin, P., Miletić, S., Stevenson, N. R., Trutti, A. C., Tse, D. H. Y., Heathcote, A., Matzke, D., Innes, R. J., Habli, S., Sokołowski, D. R., Alkemade, A., Håberg, A. K., & Forstmann, B. U. (2023). Investigating intra-individual networks of response inhibition and interference resolution using 7T MRI. NeuroImage, 119988. 10.1016/j.neuroimage.2023.119988

Isherwood, S. J. S., Kemp, S., Miletić, S., Stevenson, N., Bazin, P.-L., & Forstmann, B. U. (2023). Multi-study fMRI outlooks on subcortical BOLD responses in the stop-signal paradigm. eLife. 10.7554/eLife.88652

Isherwood, S. J. S., Keuken, M. C., Bazin, P. L., & Forstmann, B. U. (2021). Cortical and subcortical contributions to interference resolution and inhibition – An fMRI ALE meta-analysis. Neuroscience & Biobehavioral Reviews, 129, 245–260. https : //doi.org/10.1016/j.neubiorev.2021.07.021

Jeurissen, B., Leemans, A., Tournier, J.-D., Jones, D. K., & Sijbers, J. (2013). Investigating the prevalence of complex fiber configurations in white matter tissue with diffusion magnetic resonance imaging. Human Brain Mapping, 34(11), 2747–2766. 10.1002/ hbm.22099

Jeurissen, B., Tournier, J.-D., Dhollander, T., Connelly, A., & Sijbers, J. (2014). Multi-tissue constrained spherical deconvolution for improved analysis of multi-shell diffusion MRI data. NeuroImage, 103, 411–426. 10.1016/j.neuroimage.2014.07.061

Jones, D. K., & Cercignani, M. (2010). Twenty-five pitfalls in the analysis of diffusion MRI data. NMR in Biomedicine, 23(7), 803–820. 10.1002/nbm.1543

Kang, W., Wang, J., & Malvaso, A. (2022). Inhibitory control in aging: The compensation-related utilization of neural circuits hypothesis. Frontiers in Aging Neuroscience, 13. https://www.frontiersin.org/articles/10.3389/fnagi.2021.771885

Kennedy, K. M., Rodrigue, K. M., Bischof, G. N., Hebrank, A. C., Reuter-Lorenz, P. A., & Park, D. C. (2015). Age trajectories of functional activation under conditions of low and high processing demands: An adult lifespan fMRI study of the aging brain. NeuroImage, 104, 21–34. 10.1016/j.neuroimage.2014.09.056

Keuken, M. C., Bazin, P.-L., Backhouse, K., Beekhuizen, S., Himmer, L., Kandola, A., Lafeber, J. J., Prochazkova, L., Trutti, A., Schäfer, A., Turner, R., & Forstmann, B. U. (2017). Effects of aging on T1, T2*, and QSM MRI values in the subcortex. Brain Structure and Function, *222*(6), 2487–2505. 10.1007/s00429-016-1352-4

Keuken, M. C., Liebrand, L. C., Bazin, P.-L., Alkemade, A., van Berendonk, N., Groot, J. M., Isherwood, S. J. S., Kemp, S. A., Lute, N., Mulder, M. J., Trutti, A. C., Caan, M. W. A., & Forstmann, B. U. (2022). A high-resolution multi-shell 3T diffusion magnetic resonance imaging dataset as part of the Amsterdam Ultra-high field adult lifespan database (AHEAD). Data in Brief, 42, 108086. 10.1016/j.dib.2022.108086

Khattar, N., Triebswetter, C., Kiely, M., Ferrucci, L., Resnick, S. M., Spencer, R. G., & Bouhrara, M. (2021). Investigation of the association between cerebral iron content and myelin content in normative aging using quantitative magnetic resonance neuroimaging. NeuroImage, 239, 118267. 10.1016/j.neuroimage.2021.118267

King, A. V., Linke, J., Gass, A., Hennerici, M. G., Tost, H., Poupon, C., & Wessa, M. (2012). Microstructure of a three-way anatomical network predicts individual differences in response inhibition: A tractography study. NeuroImage, 59(2), 1949–1959. 10.1016/ j.neuroimage.2011.09.008

Kleerekooper, I., van Rooij, S. J. H., van den Wildenberg, W. P. M., de Leeuw, M., Kahn, R. S., & Vink, M. (2016). The effect of aging on fronto-striatal reactive and proactive inhibitory control. NeuroImage, 132, 51–58. 10.1016/j.neuroimage.2016.02.031

Lazari, A., & Lipp, I. (2021). Can MRI measure myelin? Systematic review, qualitative assessment, and meta-analysis of studies validating microstructural imaging with myelin histology. NeuroImage, 230, 117744. 10.1016/j.neuroimage.2021.117744

Lebel, C., Gee, M., Camicioli, R., Wieler, M., Martin, W., & Beaulieu, C. (2012). Diffusion tensor imaging of white matter tract evolution over the lifespan. NeuroImage, 60(1), 340–352. 10.1016/j.neuroimage.2011.11.094

Lee, H. W., Lu, M.-S., Chen, C.-Y., Muggleton, N. G., Hsu, T.-Y., & Juan, C.-H. (2016). Roles of the pre-SMA and rIFG in conditional stopping revealed by transcranial magnetic stimulation. Behavioural Brain Research, 296, 459–467. 10.1016/j.bbr.2015.08.024

Lin, M., Cai, G., Li, Y., Sun, Y., Song, Y., Cai, G., & Jiang, R. (2023). Association between beta oscillations from subthalamic nucleus and quantitative susceptibility mapping in deep gray matter structures in Parkinson’s disease. Brain Sciences, 13(1), 81. 10.3390/ brainsci13010081

Lofredi, R., Auernig, G. C., Irmen, F., Nieweler, J., Neumann, W.-J., Horn, A., Schneider, G.-H., & Kühn, A. A. (2021). Subthalamic stimulation impairs stopping of ongoing movements. Brain, 144(1), 44–52. 10.1093/brain/awaa341

Madden, D. J., & Merenstein, J. L. (2023). Quantitative susceptibility mapping of brain iron in healthy aging and cognition. NeuroImage, 282, 120401. https :// doi . org / 10 . 1016 / j . neuroimage.2023.120401

Madsen, K. S., Johansen, L. B., Thompson, W. K., Siebner, H. R., Jernigan, T. L., & Baaré, W. F. C. (2020). Maturational trajectories of white matter microstructure underlying the right presupplementary motor area reflect individual improvements in motor response can-cellation in children and adolescents. NeuroImage, 220, 117105. 10.1016/j.neuroimage.2020.117105

Marques, J. P., Kober, T., Krueger, G., van der Zwaag, W., Van de Moortele, P.-F., & Gruetter, R. (2010). MP2RAGE, a self bias-field corrected sequence for improved segmentation and T1-mapping at high field. NeuroImage, 49(2), 1271–1281. 10.1016/j.neuroimage.2009.10.002

Miletić, S., Bazin, P.-L., Isherwood, S. J. S., Keuken, M. C., Alkemade, A., & Forstmann, B. U. (2022). Charting human subcortical maturation across the adult lifespan with in vivo 7 T MRI. NeuroImage, 249, 118872. 10.1016/j.neuroimage.2022.118872

Miletić, S., Bazin, P.-L., Weiskopf, N., van der Zwaag, W., Forstmann, B. U., & Trampel, R. (2020). fMRI protocol optimization for simultaneously studying small subcortical and cortical areas at 7T. NeuroImage, 219(116992). 10.1016/j.neuroimage.2020.116992

Moreau, B., Iannessi, A., Hoog, C., & Beaumont, H. (2018). How reliable are ADC measurements? A phantom and clinical study of cervical lymph nodes. European Radiology, 28(8), 3362– 3371. 10.1007/s00330-017-5265-2

Munakata, Y., Herd, S. A., Chatham, C. H., Depue, B. E., Banich, M. T., & O’Reilly, R. C. (2011). A unified framework for inhibitory control. Trends in Cognitive Sciences, 15(10), 453–459. 10.1016/j.tics.2011.07.011

Nambu, A., Tokuno, H., & Takada, M. (2002). Functional significance of the cortico–subthalamo–pallidal ‘hyperdirect’ pathway. Neuroscience Research, 43(2), 111–117. 10.1016/ S0168-0102(02)00027-5

Narayanan, N. S., Wessel, J. R., & Greenlee, J. D. W. (2020). The fastest way to stop: Inhibitory control and IFG-STN hyperdirect connectivity. Neuron, 106(4), 549–551. 10.1016/j.neuron.2020.04.017

Neubert, F.-X., Mars, R. B., Sallet, J., & Rushworth, M. F. S. (2015). Connectivity reveals relationship of brain areas for reward-guided learning and decision making in human and monkey frontal cortex. Proceedings of the National Academy of Sciences, 112(20), E2695–E2704. 10.1073/pnas.1410767112

Neubert, F.-X., Mars, R. B., Thomas, A. G., Sallet, J., & Rushworth, M. F. (2014). Comparison of human ventral frontal cortex areas for cognitive control and language with areas in monkey frontal cortex. Neuron, 81(3), 700–713. 10.1016/j.neuron.2013.11.012

Nikitenko, T., Chowdhury, N., Puri, R., & He, J. L. (2020). Response inhibition in humans: A whistle stop review. Journal of Neurophysiology, 123(3), 861–864. 10.1152/jn.00572.2019

Nougaret, S., Meffre, J., Duclos, Y., Breysse, E., & Pelloux, Y. (2013). First evidence of a hyper-direct prefrontal pathway in the primate: Precise organization for new insights on subthalamic nucleus functions. Frontiers in Computational Neuroscience, 7, 135. 10.3389/fncom.2013.00135

O’Donnell, L. J., & Westin, C.-F. (2011). An introduction to diffusion tensor image analysis. Neurosurgery Clinics of North America, 22(2), 185–viii. 10.1016/j.nec.2010.12.004

Palmer, A. L., & Ousman, S. S. (2018). Astrocytes and aging [Publisher: Frontiers]. Frontiers in Aging Neuroscience, 10. 10.3389/fnagi.2018.00337

Park, D. C., & Reuter-Lorenz, P. (2009). The adaptive brain: Aging and neurocognitive scaffolding. Annual Review of Psychology, 60, 173–196. 10.1146/annurev.psych.59.103006.093656

Peters, A. (2009). The effects of normal aging on myelinated nerve fibers in monkey central nervous system. Frontiers in Neuroanatomy, 3. 10.3389/neuro.05.011.2009

Porcu, M., Cocco, L., Puig, J., Mannelli, L., Yang, Q., Suri, J. S., Defazio, G., & Saba, L. (2021). Global fractional anisotropy: Effect on resting-state neural activity and brain networking in healthy participants. Neuroscience, 472, 103–115. 10.1016/j.neuroscience.2021.07.021

Puri, R., Nikitenko, T., & Kemp, S. A. (2018). Using transcranial magnetic stimulation to investigate the neural mechanisms of inhibitory control. Journal of Neurophysiology, 120(4), 1587–1590. 10.1152/jn.00366.2018

Quetscher, C., Yildiz, A., Dharmadhikari, S., Glaubitz, B., Schmidt-Wilcke, T., Dydak, U., & Beste, C. (2015). Striatal GABA-MRS predicts response inhibition performance and its cortical electrophysiological correlates. Brain Structure and Function, 220(6), 3555–3564. 10.1007/s00429-014-0873-y

Radtke, C., Spies, M., Sasaki, M., Vogt, P., & Kocsis, J. D. (2007). Demyelinating diseases and potential repair strategies. International Journal of Developmental Neuroscience, 25(3), 149–153. 10.1016/j.ijdevneu.2007.02.002

Rae, C., Hughes, L., Anderson, M., & Rowe, J. (2015). The prefrontal cortex achieves inhibitory control by facilitating subcortical motor pathway connectivity. Journal of Neuroscience, 35(2), 786–794. 10.1523/JNEUROSCI.3093-13.2015

Rawlings, J. O., Pantula, S. G., & Dickey, D. A. (Eds.). (1998). Applied Regression Analysis. Springer-Verlag. 10.1007/b98890

Reuter-Lorenz, P. A., & Park, D. C. (2014). How does it STAC up? Revisiting the Scaffolding Theory of Aging and Cognition. Neuropsychology Review, 24(3), 355–370. 10.1007/s11065-014-9270-9

Rey-Mermet, A., & Gade, M. (2018). Inhibition in aging: What is preserved? What declines? A meta-analysis. Psychonomic Bulletin & Review, 25(5), 1695–1716. https : / / doi . org / 10 . 3758/s13423-017-1384-7

Rocha, G. S., Freire, M. A. M., Britto, A. M., Paiva, K. M., Oliveira, R. F., Fonseca, I. A. T., Araújo, D. P., Oliveira, L. C., Guzen, F. P., Morais, P. L. A. G., & Cavalcanti, J. R. L. P. (2023). Basal ganglia for beginners: The basic concepts you need to know and their role in movement control. Frontiers in Systems Neuroscience, 17, 1242929. 10.3389/fnsys.2023.1242929

Scheurer, E., Lovblad, K.-O., Kreis, R., Maier, S., Boesch, C., Dirnhofer, R., & Yen, K. (2011). Forensic application of postmortem diffusion-weighted and diffusion tensor MR imaging of the human brain in situ. AJNR: American Journal of Neuroradiology, 32(8), 1518–1524. 10.3174/ajnr.A2508

Schilling, K. G., Tax, C. M. W., Rheault, F., Landman, B. A., Anderson, A. W., Descoteaux, M., & Petit, L. (2021). Prevalence of white matter pathways coming into a single white matter voxel orientation: The bottleneck issue in tractography. Human Brain Mapping, 43(4), 1196–1213. 10.1002/hbm.25697

Schmidt, R., & Berke, J. D. (2017). A Pause-then-Cancel model of stopping: Evidence from basal ganglia neurophysiology. *Philosophical Transactions of the Royal Society of London. Series B*, Biological Sciences, 372(1718), 20160202. 10.1098/rstb.2016.0202

Schmidt, R., Leventhal, D. K., Mallet, N., Chen, F., & Berke, J. D. (2013). Canceling actions involves a race between basal ganglia pathways. Nature Neuroscience, 16(8), 1118–1124. 10.1038/nn.3456

Schmidt, S., Cichy, R. M., Kraft, A., Brocke, J., Irlbacher, K., & Brandt, S. A. (2009). An initial transient-state and reliable measures of corticospinal excitability in TMS studies. Clinical Neurophysiology, 120(5), 987–993. 10.1016/j.clinph.2009.02.164

Schroll, H., & Hamker, F. H. (2013). Computational models of basal-ganglia pathway functions: Focus on functional neuroanatomy [Publisher: Frontiers]. Frontiers in Systems Neuroscience, 7. 10.3389/fnsys.2013.00122

Schwarz, G. (1978). Estimating the dimension of a model. The Annals of Statistics, 6(2), 461–464. 10.1214/aos/1176344136

Seabold, S., & Perktold, J. (2010). Statsmodels: Econometric and statistical modeling with Python, 92–96. 10.25080/Majora-92bf1922-011

Sebastian, A., Baldermann, C., Feige, B., Katzev, M., Scheller, E., Hellwig, B., Lieb, K., Weiller, C., Tüscher, O., & Klöppel, S. (2013). Differential effects of age on subcomponents of response inhibition. Neurobiology of Aging, 34(9), 2183–2193. 10.1016/j.neurobiolaging.2013.03.013

Sebastian, A., Pohl, M. F., Klöppel, S., Feige, B., Lange, T., Stahl, C., Voss, A., Klauer, K. C., Lieb, K., & Tüscher, O. (2013). Disentangling common and specific neural subprocesses of response inhibition. NeuroImage, 64, 601–615. 10.1016/j.neuroimage.2012.09.020

Seiler, A., Schöngrundner, S., Stock, B., Nöth, U., Hattingen, E., Steinmetz, H., Klein, J. C., Baudrexel, S., Wagner, M., Deichmann, R., & Gracien, R.-M. (2020). Cortical aging – new insights with multiparametric quantitative MRI. Aging (Albany NY), 12(16), 16195–16210. 10.18632/aging.103629

Sener, R. N. (2001). Diffusion MRI: Apparent diffusion coefficient (ADC) values in the normal brain and a classification of brain disorders based on ADC values. Computerized Medical Imaging and Graphics, 25(4), 299–326. 10.1016/S0895-6111(00)00083-5

Sharp, D. J., Bonnelle, V., De Boissezon, X., Beckmann, C. F., James, S. G., Patel, M. C., & Mehta, M. A. (2010). Distinct frontal systems for response inhibition, attentional capture, and error processing. Proceedings of the National Academy of Sciences of the United States of America, 107(13), 6106–6111. 10.1073/pnas.1000175107

Shin, C., Lee, S., Lee, J.-Y., Rhim, J. H., & Park, S.-W. (2018). Non-motor symptom burdens are not associated with iron accumulation in early Parkinson’s disease: A quantitative susceptibility mapping study. Journal of Korean Medical Science, 33(13), e96. 10.3346/jkms.2018.33.e96

Singh, M., Fuelscher, I., He, J., Anderson, V., Silk, T. J., & Hyde, C. (2021). Inter-individual performance differences in the stop-signal task are associated with fibre-specific microstructure of the fronto-basal-ganglia circuit in healthy children. Cortex, 142, 283–295. 10.1016/j.cortex.2021.06.002

Smith, R. E., Tournier, J.-D., Calamante, F., & Connelly, A. (2015). SIFT2: Enabling dense quantitative assessment of brain white matter connectivity using streamlines tractography. NeuroImage, 119, 338–351. 10.1016/j.neuroimage.2015.06.092

Smith, S. M., Jenkinson, M., Woolrich, M. W., Beckmann, C. F., Behrens, T. E. J., Johansen-Berg, H., Bannister, P. R., De Luca, M., Drobnjak, I., Flitney, D. E., Niazy, R. K., Saunders, J., Vickers, J., Zhang, Y., De Stefano, N., Brady, J. M., & Matthews, P. M. (2004). Advances in functional and structural MR image analysis and implementation as FSL. NeuroImage, 23, S208–S219. 10.1016/j.neuroimage.2004.07.051

Smittenaar, P., Guitart-Masip, M., Lutti, A., & Dolan, R. J. (2013). Preparing for selective inhibition within frontostriatal loops [Publisher: Society for Neuroscience Section: Articles]. Journal of Neuroscience, 33(46), 18087–18097. 10.1523/JNEUROSCI.2167-13.2013

Suda, A., Osada, T., Ogawa, A., Tanaka, M., Kamagata, K., Aoki, S., Hattori, N., & Konishi, S. (2020). Functional organization for response inhibition in the right inferior frontal cortex of individual human brains. Cerebral Cortex, 30(12), 6325–6335. 10.1093/ cercor/bhaa188

Sullivan, E. V., & Pfefferbaum, A. (2006). Diffusion tensor imaging and aging. Neuroscience and Biobehavioral Reviews, 30(6), 749–761. 10.1016/j.neubiorev.2006.06.002

Sun, F., Huang, Y., Wang, J., Hong, W., & Zhao, Z. (2023). Research progress in diffusion spectrum imaging [Number: 10 Publisher: Multidisciplinary Digital Publishing Institute]. Brain Sciences, 13(10), 1497. 10.3390/brainsci13101497

Tan, E. T., Marinelli, L., Sperl, J. I., Menzel, M. I., & Hardy, C. J. (2015). Multi-directional anisotropy from diffusion orientation distribution functions. Journal of Magnetic Resonance Imaging, 41(3), 841–850. 10.1002/jmri.24589

Tardif, C. L., Gauthier, C. J., Steele, C. J., Bazin, P.-L., Schäfer, A., Schaefer, A., Turner, R., & Villringer, A. (2016). Advanced MRI techniques to improve our understanding of experience-induced neuroplasticity. NeuroImage, 131, 55–72. 10.1016/j.neuroimage.2015.08.047

Teipel, S. J., Lerche, M., Kilimann, I., O’Brien, K., Grothe, M., Meyer, P., Li, X., Sänger, P., & Hauenstein, K. (2014). Decline of fiber tract integrity over the adult age range: A diffusion spectrum imaging study [_eprint: https://onlinelibrary.wiley.com/doi/pdf/10.1002/jmri.24420]. *Journal of Magnetic Resonance Imaging*, *40*(2), 348–359. 10.1002/jmri.24420

Tennyson, S. S., Brockett, A. T., Hricz, N. W., Bryden, D. W., & Roesch, M. R. (2018). Firing of Putative Dopamine Neurons in Ventral Tegmental Area Is Modulated by Probability of Success during Performance of a Stop-Change Task. eNeuro, 5(2), ENEURO.0007– 18.2018. 10.1523/ENEURO.0007-18.2018

Tournier, J.-D., Calamante, F., & Connelly, A. (2010). Improved probabilistic streamlines tractography by 2nd order integration over fibre orientation distributions. Proceedings of the International Society for Magnetic Resonance in Medicine, 18, 1670.

Tournier, J.-D., Calamante, F., Gadian, D. G., & Connelly, A. (2004). Direct estimation of the fiber orientation density function from diffusion-weighted MRI data using spherical deconvolution. NeuroImage, 23(3), 1176–1185. 10.1016/j.neuroimage.2004.07.037

Tournier, J.-D., Smith, R., Raffelt, D., Tabbara, R., Dhollander, T., Pietsch, M., Christiaens, D., Jeurissen, B., Yeh, C.-H., & Connelly, A. (2019). MRtrix3: A fast, flexible and open software framework for medical image processing and visualisation. NeuroImage, 202, 116137. 10.1016/j.neuroimage.2019.116137

Tsvetanov, K., Ye, Z., Hughes, L., Samu, D., Treder, M., Wolpe, N., Tyler, L., Rowe, J., & Cambridge, C. f. A. N. (2018). Activity and connectivity differences underlying inhibitory control across the adult life span. Journal of Neuroscience, 38(36), 7887–7900. 10.1523/JNEUROSCI.2919-17.2018

Tubi, M. A., Feingold, F. W., Kothapalli, D., Hare, E. T., King, K. S., Thompson, P. M., Braskie, M. N., & Alzheimer’s Disease Neuroimaging Initiative. (2020). White matter hyperintensities and their relationship to cognition: Effects of segmentation algorithm. NeuroImage, *206*, 116327. 10.1016/j.neuroimage.2019.116327

Tuch, D. S. (2004). Q-ball imaging [_eprint: https://onlinelibrary.wiley.com/doi/pdf/10.1002/mrm.20279]. *Magnetic Resonance in Medicine*, *52*(6), 1358–1372. 10.1002/mrm.20279

van de Laar, M., Van Den Wildenberg, W., van Boxtel, G., & van der Molen, M. (2011). Lifespan changes in global and selective stopping and performance adjustments. Frontiers in Psychology, 2. 10.3389/fpsyg.2011.00357

Verbruggen, F., Aron, A. R., Band, G. P., Beste, C., Bissett, P. G., Brockett, A. T., Brown, J. W., Chamberlain, S. R., Chambers, C. D., Colonius, H., Colzato, L. S., Corneil, B. D., Coxon, J. P., Dupuis, A., Eagle, D. M., Garavan, H., Greenhouse, I., Heathcote, A., Huster, R. J., . . . Boehler, C. N. (2019). A consensus guide to capturing the ability to inhibit actions and impulsive behaviors in the stop-signal task. eLife, 8, e46323. 10.7554/eLife.46323

Verstraelen, S., Cuypers, K., Maes, C., Hehl, M., Van Malderen, S., Levin, O., Mikkelsen, M., Meesen, R., & Swinnen, S. (2021). Neurophysiological modulations in the (pre)motor-motor network underlying age-related increases in reaction time and the role of GABA levels – a bimodal TMS-MRS study. NeuroImage, 243. 10.1016/j.neuroimage.2021.118500

Volz, L. J., Cieslak, M., & Grafton, S. T. (2018). A probabilistic atlas of fiber crossings for variability reduction of anisotropy measures. Brain Structure and Function, 223(2), 635–651. 10.1007/s00429-017-1508-x

Weiskopf, N., Mohammadi, S., Lutti, A., & Callaghan, M. F. (2015). Advances in MRI-based computational neuroanatomy: From morphometry to in-vivo histology. Current Opinion in Neurology, 28(4), 313–322. 10.1097/WCO.0000000000000222

Wessel, J. R. (2020). Beta-bursts reveal the trial-to-trial dynamics of movement initiation and cancellation. Journal of Neuroscience, 40(2), 411–423. 10.1523/JNEUROSCI.1887-19.2019

Wessel, J. R., & Aron, A. R. (2017). On the globality of motor suppression: Unexpected events and their influence on behavior and cognition. Neuron, 93(2), 259–280. 10.1016/j.neuron.2016.12.013

Williams, B. R., Ponesse, J. S., Schachar, R. J., Logan, G. D., & Tannock, R. (1999). Development of inhibitory control across the life span. Developmental Psychology, 35(1), 205–213. 10.1037//0012-1649.35.1.205

Williamson, J. M., & Lyons, D. A. (2018). Myelin dynamics throughout life: An ever-changing landscape? Frontiers in Cellular Neuroscience, 12. 10.3389/fncel.2018.00424

Xu, B., Sandrini, M., Wang, W.-T., Smith, J. F., Sarlls, J. E., Awosika, O., Butman, J. A., Horwitz, B., & Cohen, L. G. (2016). PreSMA stimulation changes task-free functional connectivity in the fronto-basal-ganglia that correlates with response inhibition efficiency. Human Brain Mapping, 37(9), 3236–3249. 10.1002/hbm.23236

Yang, M.-H., Yao, Z.-F., & Hsieh, S. (2019). Multimodal neuroimaging analysis reveals age-associated common and discrete cognitive control constructs. Human Brain Mapping, 40(9), 2639–2661. 10.1002/hbm.24550

Yeatman, J. D., Wandell, B. A., & Mezer, A. A. (2014). Lifespan maturation and degeneration of human brain white matter. Nature Communications, 5(1), 4932. 10.1038/ ncomms5932

Zandbelt, B. B., Bloemendaal, M., Hoogendam, J. M., Kahn, R. S., & Vink, M. (2013). Transcranial magnetic stimulation and functional MRI reveal cortical and subcortical interactions during stop-signal response inhibition. Journal of Cognitive Neuroscience, 25(2), 157–174. 10.1162/jocn_a_00309

Zhang, F., & Iwaki, S. (2020). Correspondence between effective connections in the stop-signal task and microstructural correlations. Frontiers in Human Neuroscience, 14, 279. https ://doi.org/10.3389/fnhum.2020.00279

Zhang, X., Huang, N., Xiao, L., Wang, F., & Li, T. (2021). Replenishing the aged brains: Targeting oligodendrocytes and myelination? Frontiers in Aging Neuroscience, 13. 10.3389/fnagi.2021.760200

Zhu, D. C., Zacks, R. T., & Slade, J. M. (2010). Brain activation during interference resolution in young and older adults: An fMRI study. NeuroImage, 50(2), 810–817. 10.1016/j.neuroimage.2009.12.087

Zimmermann, J., Ritter, P., Shen, K., Rothmeier, S., Schirner, M., & McIntosh, A. R. (2016). Structural architecture supports functional organization in the human aging brain at a regionwise and network level. Human Brain Mapping, 37(7), 2645–2661. 10.1002/hbm.23200

