## Supplementary Materials for "The ageing stopping network: Regional and network changes in the IFG, preSMA, and STN across the adult lifespan"

### From 3.1: *t*-test results

Results of the Student *t*-tests, assessing if there were any significant differences between hemispheres for any of the measures.

Table S1: Student *t*-test results

| Measure | Student's *t* | *p*-value |
|---|---|---|
| Regional iron | -0.35 | .730 |
| Tract iron | 0.69 | .493 |
| Regional myelin | -0.21 | .832 |
| Tract myelin | 0.13 | .897 |
| GFA | 0.83 | .408 |
| ADC | -0.69 | .494 |

Given all $p > .05$, we collapsed across hemispheres for all analyses to increase statistical power.

### From 3.2 and 3.3.1: Model candidates

The following models were considered as potential candidates, with $\gamma$ denoting the outcome variable (iron and myelin in 3.2; iron, myelin, GFA, and ADC in 3.3.1):

$$\gamma_1 = \beta_1 \text{Age}$$

$$\gamma_2 = \beta_1 \text{Age} + \beta_2 \text{Age}^2$$

$$\gamma_3 = \beta_1 \text{Age} + \beta_2 \text{Age}^2 + \beta_3 \text{Age}^3$$

### From 3.3.2: Mediation results

Results from the mediation analysis. Statistically significant results are emphasised with *



PREPRINT

| DWI metric | Tract | Mediator | Analysis | Estimate | *p*-value |
|---|---|---|---|---|---|
| ADC | IFG-preSMA | Iron | ACME | -0.042 | .030* |
| | | | ADE | 0.322 | <.001* |
| | | | Total | 0.281 | <.001* |
| | | Myelin | ACME | -0.015 | .414 |
| | | | ADE | 0.262 | <.001* |
| | | | Total | 0.247 | <.001* |
| | STN-IFG | Iron | ACME | -0.019 | .212 |
| | | | ADE | 0.262 | <.001* |
| | | | Total | 0.243 | .002* |
| | | Myelin | ACME | -0.021 | .184 |
| | | | ADE | 0.208 | <.001* |
| | | | Total | 0.187 | .010* |
| | STN-preSMA | Iron | ACME | -0.028 | .080 |
| | | | ADE | 0.259 | <.001* |
| | | | Total | 0.231 | <.001* |
| | | Myelin | ACME | -0.016 | .282 |
| | | | ADE | 0.211 | <.001* |
| | | | Total | 0.194 | <.001* |
| GFA | IFG-preSMA | Iron | ACME | -0.013 | .886 |
| | | | ADE | 0.148 | .334 |
| | | | Total | 0.134 | .420 |
| | | Myelin | ACME | -0.170 | .832 |
| | | | ADE | 0.093 | .526 |
| | | | Total | 0.076 | .640 |
| | STN-IFG | Iron | ACME | -0.045 | .642 |
| | | | ADE | 0.320 | .018* |
| | | | Total | 0.275 | .080 |
| | | Myelin | ACME | 0.001 | .956 |
| | | | ADE | 0.104 | .416 |
| | | | Total | 0.104 | .442 |
| | STN-preSMA | Iron | ACME | -0.048 | .550 |
| | | | ADE | 0.273 | .036* |
| | | | Total | 0.225 | .124 |
| | | Myelin | ACME | -0.015 | .760 |
| | | | ADE | 0.141 | .280 |
| | | | Total | 0.126 | .368 |

PREPRINT
